# Hemagglutinin of Influenza A, but not of Influenza B and C viruses is acylated by ZDHHC2, 8, 15 and 20

**DOI:** 10.1101/2019.12.15.877001

**Authors:** Mohamed Rasheed Gadalla, Laurence Abrami, F Gisou van der Goot, Michael Veit

## Abstract

Hemagglutinin (HA), a glycoprotein of Influenza A viruses and its proton-channel M2 are site-specifically modified with fatty acids. Whereas two cysteines in the short cytoplasmic tail of HA contain only palmitate, stearate is exclusively attached to one cysteine located at the cytoplasmic border of the transmembrane region (TMR). M2 is palmitoylated at a cysteine positioned in an amphiphilic helix near the TMR. The enzymes catalyzing acylation of HA and M2 have not been identified, but zinc finger DHHC domain containing (ZDHHC) palmitoyltransferases are candidates. We used a siRNA library to knockdown expression of each of the 23 human ZDHHCs in HA-expressing HeLa cells. siRNAs against ZDHHC2 and 8 had the strongest effect on acylation of HA as demonstrated by acyl-RAC and confirmed by 3H-palmitate labelling. CRISPR/Cas9 knockout of ZDHHC2 and 8 in HAP1 cells, but also of the phylogenetically related ZDHHCs 15 and 20 strongly reduced acylation of group 1 and group 2 HAs and of M2, but individual ZDHHCs exhibit slightly different substrate preferences. These ZDHHCs co- localize with HA at membranes of the exocytic pathway in a human lung cell line. ZDHHC2, 8, 15 and 20 are not required for acylation of the hemagglutinin-esterase-fusion protein of Influenza C virus that contains only stearate at one transmembrane cysteine. Knockout of these ZDHHCs also did not compromise acylation of HA of Influenza B virus that contains two palmitoylated cysteines in its cytoplasmic tail. Results are discussed with respect to the acyl preferences and possible substrate recognition features of the identified ZDHHCs.

## 1. Introduction

In influenza A and B viruses there are two viral spikes embedded in the envelope: the trimeric hemagglutinin (HA), which catalyzes virus entry by binding to sialic acid containing receptors and by performing fusion of viral with endosomal membranes and the neuraminidase (NA), which is required for release of virus particles from infected cells [1]. In Influenza C virus all three activities (Receptor binding and destroying, membrane fusion) are combined in one spike, which is designated hemagglutinin-esterase-fusion (HEF) [2]. Influenza A virus particles contain also minor amounts of the tetrameric M2 proton channel. M2 conducts protons from the acidic milieu of the endosome into virus particles, which catalyzes uncoating of the viral genome. In addition, an amphiphilic helix located adjacent to the transmembrane region is required for virus budding [3].

HA and HEF as well as M2 are S-acylated at cytoplasmic and transmembrane cysteine residues, but with different acyl chains [4-9]. HA of Influenza B virus contains 97% palmitate attached to two cytoplasmic cysteine residues and HEF of Influenza C virus is predominantly (88%) stearoylated at one cysteine at the end of the transmembrane region (TMR). HAs of Influenza A virus contain a mixture of palmitate and stearate, but MS analysis revealed that stearate is exclusively attached to the cysteine positioned at the end of the transmembrane region, whereas the two cytoplasmic cysteines are always acylated with palmitate, attachment of stearate was always less than 5% [10, 11]. The percentage of stearate in all HA variants differs from 35% (meaning that each TMR cysteine contains stearate) to 12% (indicating that only one out of three TMR cysteines in the trimeric HA spike is stearoylated). However, the main signal for stearate attachment is the location of an acylation site relative to the transmembrane span since shifting the TMR cysteine to a cytoplasmic location virtually eliminated attachment of stearate [12].

Inconsistent data have been published whether acylation of HA is required for cell entry of viruses via membrane fusion [13], but the acyl chains are necessary for incorporation of HA into nanodomains of the plasma membrane [14, 15] and for incorporation of M1 into virus particles [16, 17]. Whatever the exact role of HA’s acylation might be, the modification is essential for virus replication since nobody could generate Influenza virus mutants of various subtypes with two or three HA acylation sites removed and mutants with one deleted site revealed greatly reduced growth rates in cell culture [16, 18, 19]. In general, removal of palmitoylated cysteine residues had a more drastic effect on virus replication than deletion of the stearate attachment site. Likewise, removing the stearoylated cysteine from the HEF protein of Influenza C virus generated a virus that was only slightly compromised in growth [20]. Reverse genetics studies have been not performed with Influenza B virus, but exchange of the conserved acylation sites in expressed HA inhibited fusion pore formation suggesting that acylation significantly contributes to virus replication [21, 22]. In summary, palmitoylation of the two cytoplasmic cysteines is essential for virus replication, whereas the stearoylated cysteine has only a modest effect.

This is supported by a comparison of all Flu A HA sequences present in the NCBI-database (≈ 17.000). Each molecule contains three, some even four cysteine residues located in the relevant region [23]. The conservation of cysteines in a highly variable molecule like HA reinforces the notion that they play an essential role for the life cycle of Influenza viruses. In contrast, the acylation site of M2 is not present in ∼15% of M2 variants and virus mutants with deleted acylation site revealed no growth defect in cell culture [24], but acylation of M2 and HA synergistically affect virus release [25] and virus mutants were attenuated in mice [26]. Thus, inhibiting the enzyme that attaches palmitate to Influenza virus proteins might be a promising strategy to block virus replication.

Enzymes that acylate HA (or any other viral glycoprotein) have not been identified, but cellular proteins are acylated by members of the ZDHHC family of proteins [27-30]. ZDHHC proteins are polytopic membrane proteins containing an Asp-His-His-Cys (DHHC) motif as the catalytic centre in one of their cytoplasmic domains. The DHHC-motif is embedded within a cysteine-rich domain (CRD) that is a variant of the zinc finger motif [31]. Most ZDHHCs exhibit a two-step reaction mechanism; a fatty acid is first transferred from the lipid donor acyl-CoA to the cysteine of the DHHC motif (Autoacylation) and subsequently to a cysteine of the substrate protein [32]. A multitude of ZDHHC proteins exists in eukaryotic cells, seven in yeast and 23 in humans. Besides the cysteine-rich domain, little sequence conservation occurs between ZDHHC proteins. Their size varies from 263 to 765 amino acids and the number of (predicted) transmembrane regions from four to six. Most ZDHHC proteins are abundant in many tissues, but some are expressed in only a few cell types. The majority of ZDHHC proteins localize to endoplasmic reticulum (ER) or Golgi membranes, and a small number target to the plasma membrane [33].

A multitude of studies showed that many cellular proteins could be palmitoylated by several, but not each of the various ZDHHC proteins indicating that the 23 enzymes show distinct, but overlapping substrate specificities [28, 34, 35]. If only a few ZDHHC proteins are involved in acylation of HA they might be promising drug targets since their blockade will result in suppression of virus replication, while acylation of cellular proteins might not be (or very little) compromised [25]. However, essentially nothing is known about how ZDHHC proteins recognize transmembrane proteins. Since HA and HEF are typical type 1 transmembrane proteins having only a very short cytoplasmic tail (3-11 amino acids) the amino acids that could interact with a ZDHHC must be present in small and well-defined region. M2 also has only one transmembrane region; the fatty acid is attached to the beginning of the amphiphilic helix adjacent to the transmembrane region. Furthermore, 3D structures of the M2 proton channel and parts of the transmembrane region of HA are available [36, 37]. Thus, identifying the ZDHHC relevant for acylation of HA and M2 might give a hint which feature in a transmembrane protein is recognized by a certain ZDHHC protein.

Likewise, only some ZDHHCs have been characterized with respect to their lipid substrate specificity. Using purified ZDHHCs and substrate proteins it was shown that ZDHHC3 has a strong preference for Pal-CoA (C16:0) over Stear-CoA (C18:0), whereas ZDHHC2 displays no clear preference [32]. Using labelling of cells with various fatty acid analogues and subsequent click-chemistry based detection it was reported that ZDHHC3, 5, 7, 11, and 15 prefer Myr-CoA (C14:0) and Pal-CoA (C16:0) over C18:0; ZDHHC2 and 4 display no clear fatty acid preference; ZDHHC17 prefers C16:0/C18:0 over C14:0; and ZDHHC23 exhibits a strong preference for C18:0 [38]. In an autoacylation assay, ZDHHC20 prefers palmitate to myristate and stearate [39].

A molecular explanation for the lipid preference was provided by the recently determined crystal structures of human ZDHHC 20 [39]. The four transmembrane helices form a tent-like structure with the DHHC-motif located at the membrane-cytosol interface. The active site has a catalytic-triad like arrangement of aspartic acid and histidine that activate the nucleophile cysteine. The highly conserved cysteine-rich region forms six β-sheets that coordinate two zinc-ions, which impart structural stability. This part of the molecule also contains a patch of positively charged residues that bind the negatively charged phosphates of the CoA moiety. The structure of an acylated enzyme intermediate revealed that the fatty acid is inserted into a hydrophobic cavity formed by all four transmembrane regions. At the narrow end of the cavity, two amino acids located on TMR 1 and TMR 3 form a hydrogen bond, which effectively closes the groove. Most ZDHHCs contain either two bulky residues, one bulky and one small or two small amino acids at the homologues position. Thus, the presence of certain amino acids (large or small) at the end of the hydrophobic cavity regulate its deepness and determines the lipid binding specificity of a certain ZDHHC protein, i. e. the presence of two large and bulky residues limits the use of longer acyl-chains, such as stearate. Here we used siRNA screens and CRISPR/Cas9 technology to identify the ZDHHC proteins that are involved in S-acylation of Influenza virus proteins.

## 2. Materials and Methods

### 2.1 Cell lines and viruses

HeLa cells were grown in complete modified Eagle’s medium (MEM, Sigma) supplemented with 10% fetal bovine serum (FBS), 2mM L-glutamine and penicillin/streptomycin antibacterial reagent at 37°C. HAP1, A549 and MDCK cells were grown in Dulbecco’s modified Eagle’s medium (DMEM; PAN-Biotech) supplemented with 10 % heat-inactivated fetal bovine serum at 37°C with 5 % CO2. CRISPR/Cas9 knockout HAP1 cell lines with frameshift mutations in coding exon of ZDHHC1, 2, 8, 15 and 20 were purchased from Horizon Genomics (Vienna, Austria).

The following viruses were used: Human Influenza A virus A/WSN/33 (H1N1); a variant of avian fowl plaque virus A/chicken/Rostock/8/1934 (H7N1) having a mutant HA with a monobasic cleavage site [40], human Influenza B virus B/Lee/40 and human Influenza C virus (C/JHB/1/66). Influenza A and B viruses were propagated and titrated in MDCK-II cells, whereas Influenza C was propagated in MDCK-I cells. HAP1 cells were infected with influenza viruses at a multiplicity of infection (MOI) of 1. After 1h adsorption, the inoculation medium was replaced by pre-warmed medium supplemented with 0.1 % BSA and incubated for 16 hours at 37°C (for influenza A) or 33°C (for influenza B and C viruses).

### 2.2 Plasmids and antibodies

Influenza virus hemagglutinin (HA) with monobasic cleavage site from H7N1 strain was cloned in pCAGGS expression plasmid. HA tagged expression vectors of mouse ZDHHCs (2,8,15 and 20) were kindly provided by Masaki Fukata (National Institutes for Physiological Sciences, Japan[41]). Antisera against FPV and Influenza C virus were generated in rabbits. Viruses were purified from embryonated eggs (Flu C) or MDCK II cells (FPV). To generate the antiserum against the HA2 subunit of FPV purified virus was subjected to SDS-PAGE and the Coomassie-stained HA2 band was cut from the gel and used for immunization.

### 2.3 PCR, Sequencing and real-time PCR

Total genomic DNA was isolated from HAP1 cells using Invisorb Spin Tissue Mini Kit following manufacturer’s instructions (Stratec). PCR was carried out using specific primers flanking the Cas9 target site of each ZDHHC gene (Table S.1). PCR amplifications were performed using Q5 High fidelity DNA polymerase (New England Biolabs) under the following conditions: 98°C for 30s, 35 cycles of 98°C for 10s, 61-67°C (according to target gene) for 30s and 72°C for 45s with a final extension step of 72 °C for 5 min. Amplified PCR products were analyzed by electrophoresis, gel purified using GF-1 ambiclean gel extraction kit (Vivantis) and sequenced (LGC Genomics).

For real-time PCR, RNA was extracted using the RNeasy kit (Qiagen). 1µg of the total RNA extracted was reverse transcribed using random hexamers and superscript II (Invitrogen). 100ng of the cDNA was used to perform the real-time PCR using Sybr green reagent (Roche). The primers for real-time PCR in HeLa and HAP1 cells are described in [42]; the primers for real-time PCR of A549 are listed in supplementary table 2. mRNA levels were normalized using the housekeeping genes: TATA-binding protein, β-glucoronidase (in HeLa and HAP1 cells) and GAPDH in A549 cells.

### 2.4 siRNA knockdown experiments

HeLa cells (2×10^6^) were transfected with 100 pmol of validated siRNAs against all human ZDHHCs [42] (Qiagen) using interferrin transfection reagent (Polyplus). Cells were again transfected 48h later with a plasmid encoding HA of the FPV mutant. 24 hours later cells were washed with cold PBS and acylation of viral HA was assessed using Acyl-RAC procedures.

### 2.5 Acyl-Resin assisted capture (Acyl-RAC)

Protein S-acylation was analyzed by the Acyl-RAC assay as described [43], with some modifications. Transfected or infected cells in a six well plate was washed with PBS, lysed in 500µl buffer A (0.5% Triton-X100, 25 mM HEPES (pH 7.4), 25 mM NaCl, 1 mM EDTA, and protease inhibitor cocktail). Disulfide bonds were reduced by adding Tris (2-carboxyethyl) phosphin (TCEP, Carl Roth, HN95.2) to a final concentration of 10mM and incubated at RT for 30min. Free SH-groups were blocked by adding methyl methanethiosulfonate (MMTS, Sigma, 208795, dissolved in 100 mM HEPES, 1 mM EDTA, 87.5 mM SDS) to a final concentration of 1.5% (v/v) and incubated for 4 h at 40 °C. Subsequently, 3 volumes of ice-cold 100% acetone was added to the cell lysate and incubated at −20°C overnight. Precipitated proteins were pelleted at 5,000xg for 10 minutes at 4°C. Pelleted proteins were washed five times with 70% (v/v) acetone, air-dried, and then re-suspended in binding buffer (100 mM HEPES, 1 mM EDTA, 35 mM SDS). 20-30 µl of the sample was removed to check for total protein expression by western blotting. The remaining lysate was divided into two equal aliquots. One aliquot was treated with hydroxylamine (0.5 M final concentration, added from a 2M hydroxylamine stock adjusted to pH 7.4) to cleave thioester bonds. The second aliquot was treated with 0.5M Tris-HCl pH 7.4. 30µl thiopropyl Sepharose beads (Sigma, T8387), which were beforehand activated by incubation for 15 min in aqua dest, were added at the same time to capture free SH-groups. Samples were incubated with beads overnight at room temperature on a rotating wheel. The beads were then washed 5x in binding buffer and bound proteins were eluted from the beads with 2x non-reducing SDS-PAGE sample buffer for 5 minutes at 95°C. Samples were then subjected to SDS-PAGE and western blotting.

### 2.6 SDS-PAGE and western blot

After sodium dodecyl sulfate-polyacrylamide gel electrophoresis (SDS-PAGE) using 12% polyacrylamide, gels were electrophoretically transferred (200mA for 1hr) onto polyvinylidene difluoride (PVDF) membranes (GE Healthcare). After blocking (blocking solution: 5% skim milk powder in PBS with 0.1% Tween-20 (PBST)) for 1h at room temperature, membranes were incubated with the primary antibody overnight at 4°C. The following commercial antibodies were used: Influenza A virus H1N1 HA antibody from rabbit (Genetex, GTX127357, 1:3000 dilution in 5% non-fat milk), Caveolin-1(N-20) antibody from rabbit (Santa Cruz-894, 1:1000) and anti-Influenza A Virus M2 protein monoclonal antibody 14C2 from mice (abcam, ab5416, 1:2000). Monoclonal antibodies against HA of Influenza B virus (a 1:1 mixture of clones 6D12 and 1B5m, 1:1000 dilution) were kindly provided by Dr. Florian Krammer [44]. To detect H7 subtype HA we used our antiserum against the HA2 subunit of the FPV strain and to detect HEF our antiserum against Influenza C virus was used, both at a dilution of 1:3000.

After washing 3x for 10min with PBST, membranes were incubated with horseradish peroxidase-coupled secondary antibody, either recombinant anti-rabbit IgG VHH Single Domain (1:5000, abcam, ab191866) or goat anti-mouse IgG (H + L), (1:2000, Biorad, 1706516) for 1 hour at room temperature. After washing three times, signals were detected by chemiluminescence using the Pierce ECLplus reagent (Thermofisher, # 32132) and a Fusion SL camera system (Peqlab, Erlangen, Germany). The density of bands was quantified with Image J software.

### 2.7 Isotope labelling and immunoprecipitation

For [^3^H]-palmitate labelling, HeLa cells were transfected with siRNAs to knockdown ZDHHCs 1, 2, 8, 15 and 20 as well as with a scrambled siRNA. 48 hours later cells were transfected with a plasmid encoding HA of the FPV mutant using Fugene (Promega). 24 hours later cells were labelled for 2 hr with 200 µCi/ml [^3^H]-palmitic acid (9,10-3H(N)) (American Radiolabeled Chemicals, Inc.) in IM (Glasgow minimal essential medium buffered with 10 mM HEPES, pH 7.4). For immunoprecipitation (IP), cells were washed 3x in cold PBS, lysed for 30 min at 4°C in IP Buffer (0.5% Nonidet P-40, 500 mM Tris pH 7.4, 20 mM EDTA, 10 mM NaF, 2 mM benzamidin and protease inhibitor cocktail (Roche)), and insoluble material was pelleted for 3 min at 5000 rpm. Supernatants were precleared using 20µl of protein G sepahrose beads (GE Healthcare, 7-0618-01) for 1hr at 4°C. Cleared supernatants were incubated with 5µl antiserum against the HA2 subunit of HA from FPV and with 30µl protein G beads overnight at 4°C on a rotating wheel. Sepharose beads were washed 3x with IP buffer and proteins were eluted in 4x non-reducing SDS-PAGE sample buffer, boiled at 95°C for 5min and subjected to SDS-PAGE. Gels were treated with Amersham Amplify (GE Healthcare) according to manufactures instructions, dried and exposed to X-ray film.

### 2.8 Confocal microscopy

A549 cells were seeded at 50% confluency one day before transfection on glass coverslips in 24-well cell culture plates. Cells were co-transfected with plasmids (1µg each) encoding HA of FPV along with individual mouse ZDHHCs (2, 8, 15 and 20), each fused to a C-terminal HA-tag using lipofectamine 3000 transfection reagent (ThermoFisher, L3000015) according to manufacturer instructions. 24h post transfection, cells were fixed with paraformaldehyde (4% in aqua dest.) for 20min at room temperature (RT), permeabilized with Triton X-100 (0.1% in PBS) for 10 min under gentle shaking and blocked with bovine serum albumin (BSA, 1% in PBS) for 45 min at RT. Cells were stained simultaneously with antiserum against influenza virus FPV (rabbit, 1:3000 diluted in 1% BSA) and mouse monoclonal anti-HA (6E2) tag antibody (1:1000 dilution in 3% bovine serum albumin, Cell signaling, #2367) for 1hr at RT. The HA-tag antibody recognizes the amino acid sequence YPYDVPDYA which is not present in the H7 subtype HA. To further exclude any possibility of cross reactivity of our used FPV antiserum with HA tagged proteins. Cells transfected only with GST (HA tag vector control) and stained with both antibodies was used as a negative control. Afterwards, cells were washed 3x for 5 min with PBS and then treated for 1hr protected from light at RT simultaneously with fluorescent secondary antibodies, anti-rabbit Alexa flour 488 from goat (ThermoFisher, #A-11034) and anti-mouse antibody coupled to Alexa Fluor 568 (ThermoFisher, #A-11004), both at a dilution of 1:1000. Nuclei were then stained with DAPI (4’,6- diamidino-2-phenylindole, dihydrochloride, ThermoFisher, 62247, 1:1000 dilution in washing buffer) for 5 min on a shaker at RT. Cells were then washed 3 times and coverslips were mounted on glass slides with ProLong Gold antifade mounting buffer (Thermo Fisher, P36930) and allowed to cure in a dark place overnight. Cells were illuminated via laser lines at 488 nm (Alex Fluor 488) and 561 nm (Alexa Fluor 568) and visualized with the VisiScope confocal FRAP System (VisiTron Systems GmbH), equipped with iXon Ultra 888 EMCCD camera, using 60X objective. The images were then processed using ImageJ (https://imagej.nih.gov/ij/).

### 2.9 Models of ZDHHC proteins, HA and M2

Figures were created with PyMol (Molecular Graphics System, Version 2.0 Schrödinger, LLC, https://pymol.org/2/). For visualization of ZDHHC20, the pdb file 6BML was used. The hydrophobic amino acid contacting the acyl chain are described in (36) and the amino acid sequence alignment in the same reference was used to find amino acids in equivalent positions in human ZDHHC2 and ZDHHC15. The model of ZDHHC8 was created with SWISS-model (62) using the first 275 amino acids of human ZDHHC8 as target sequence. SWISS-model found the human ZDHHC20 structure (pdb file 6bmm.1. A) as the best template for modeling. The target sequence of ZDHHC8 and ZDHHC20 exhibit a sequence identity of 27.20% and a sequence similarity of 35%. Although the global QMEAN, (Qualitative Model Energy Analysis), is quite low (−5.17) the local absolute quality estimates of the four transmembrane regions is much better (inset in Fig. 8). The model of HA was created from pdb file 6HJ0, which contains the Cryo-EM structure of a full-length group 1 HA from the virus A/duck/Alberta/35/76 (H1N1) embedded in a detergent micelle (33). The model of M2 was created from pdb file 2L0J, which contains the solid-state NMR structure of amino acids 22-62 of M2 in a lipid bilayer (34).

**Figure 1:**
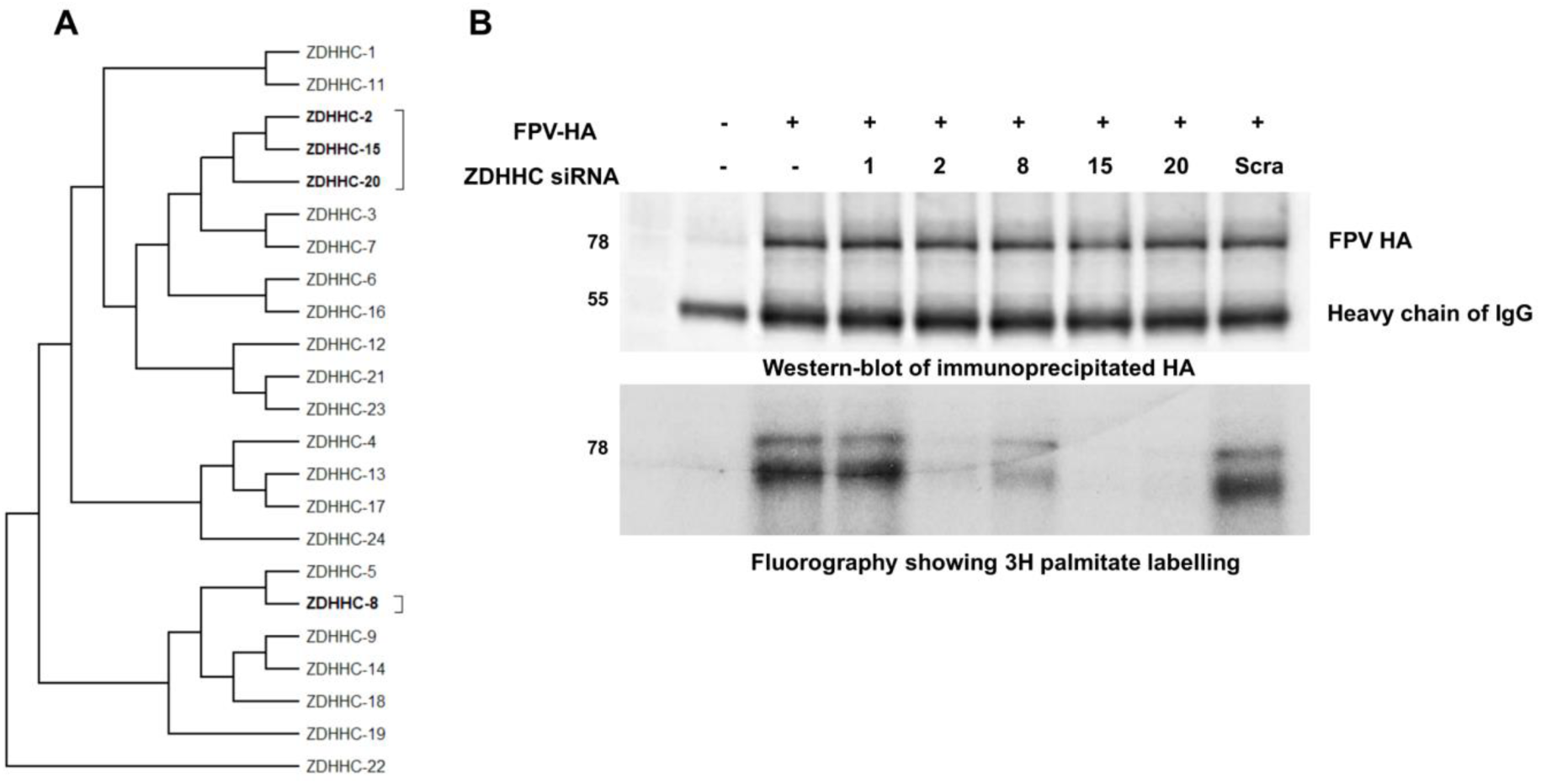
siRNAs against ZDHHC2, 8, 15 and 20 block acylation of HA as demonstrated by [^3^H] palmitate labelling. **A:** Phylogenetic tree of human ZDHHCs showing that ZDHHC2, 15 and 20 as well as 5 and 8 belong to the same group. **B:** [^3^H]-palmitate labelling: HeLa cells were transfected with siRNAs against ZDHHC1, 2, 8, 15, with a scrambled siRNA (scra) or remain un-transfected (-) and 48h later with a plasmid encoding HA from H7 subtype from FPV. After 24 hours, cells were labeled for 2 hours with [^3^H]-palmitate, lysed and subjected to immunoprecipitation with antiserum against the HA2 subunit (lower panel). An aliquot of the sample was subjected to western blotting with the same antibody (upper panel). The lower band in the upper panel is the heavy chain of the antibodies used for immuno-precipitation.

**Figure 2:**
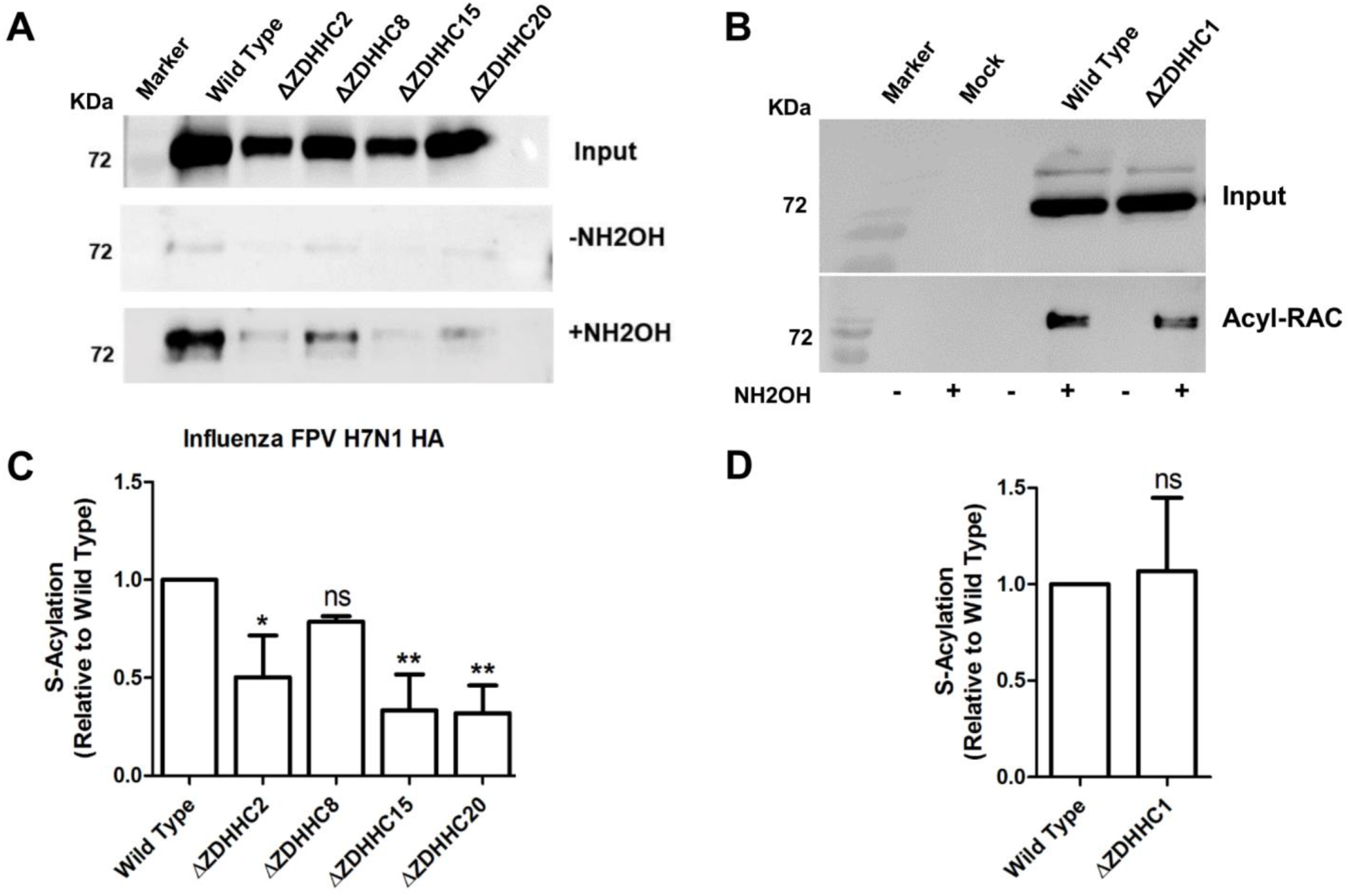
HAP1 cells deficient in ZDHHC2, 8, 15 and 20 show compromised acylation of HA. **A+B:** HAP1 cells where the indicated ZDHHCs were knocked-out using the CRISPR/Cas9 technology were infected with the variant of FPV at MOI of 1. 24 hours later, acylation of HA was analyzed using Acyl-RAC. **C+D:** Quantification of the result from this and two other identical experiments. The optical density of the +NH2OH bands was divided by the density of bands from the input and normalized to wild type (=1). The mean ± standard deviation is shown. The asterisks indicate statistically significant differences (*P < 0.05, **P < 0.01) between WT and the deficient cells. One-way ANOVA followed by Tukey’s multiple comparison test was applied for statistical analysis. ns: non-significant.

**Figure 3:**
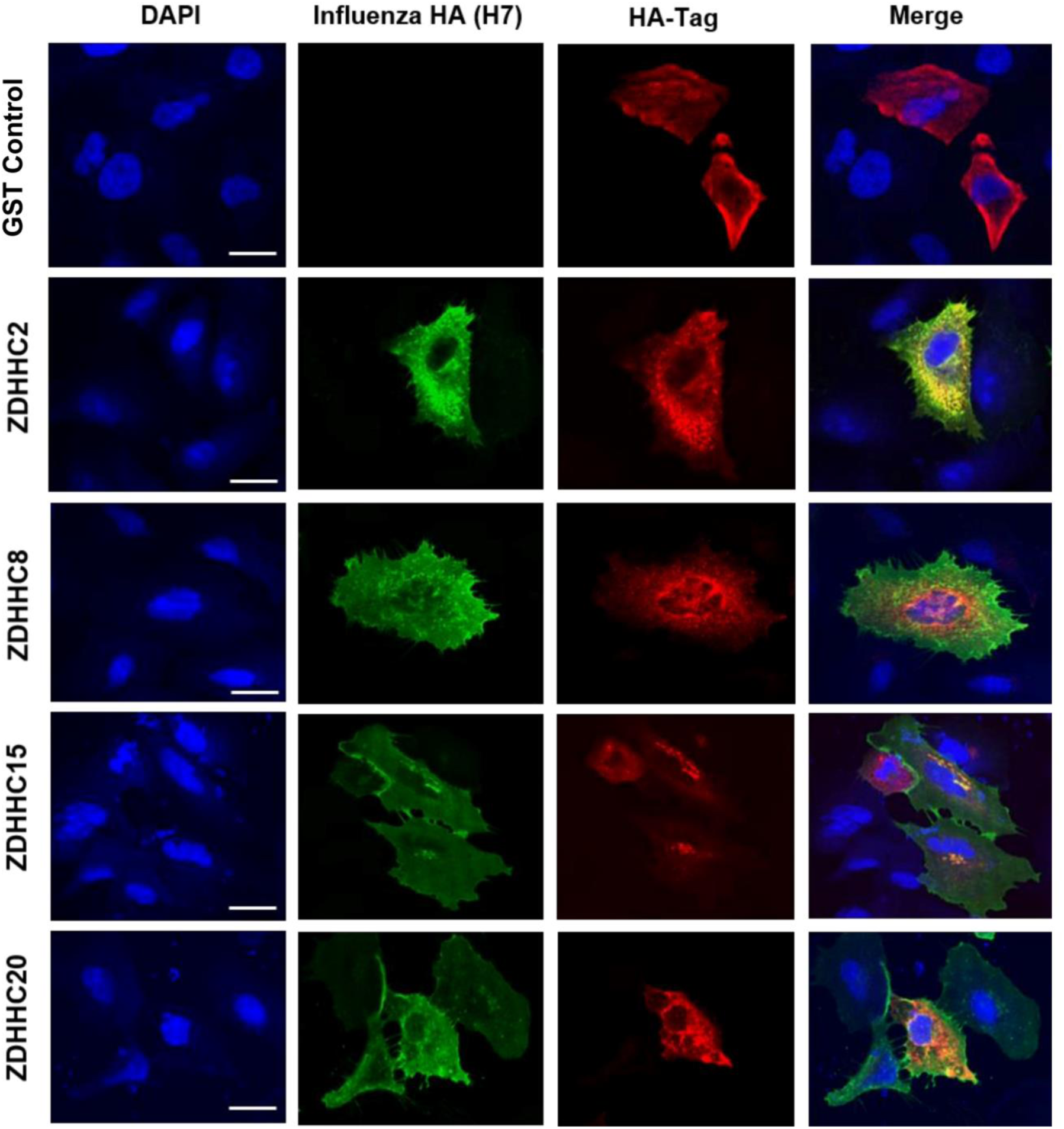
ZDHHCs 2, 8, 15 and 20 co-localize with HA at membranes of the exocytic pathway in human lung cells. A549 human lung cells were transfected with plasmids encoding H7 subtype HA and the indicated ZDHHCs fused to a C-terminal HA-tag. 24 hours later cells were fixed, permeabilized and stained with anti-FPV antiserum, and anti-HA-tag antibodies followed by secondary antibody coupled to Alexa-Fluor 568 (red for ZDHHCs) and Alexa Fluor 488 (green for HA), respectively. Nuclei were stained with DAPI. Scale bar =50µm. Cells transfected with a plasmid expressing GST and stained with both antibodies was used as a negative control. Co-localization of HA with each ZDHHC from at least 40 cells was quantified with the Pearson’s correlation coefficient method using the JACoP plugin of the ImageJ software. 60% of HA pixels overlapped with ZDHHC8, 72% with ZDHHC2, 74% with ZDHHC20 and 75% with ZDHHC15.

**Figure 4:**
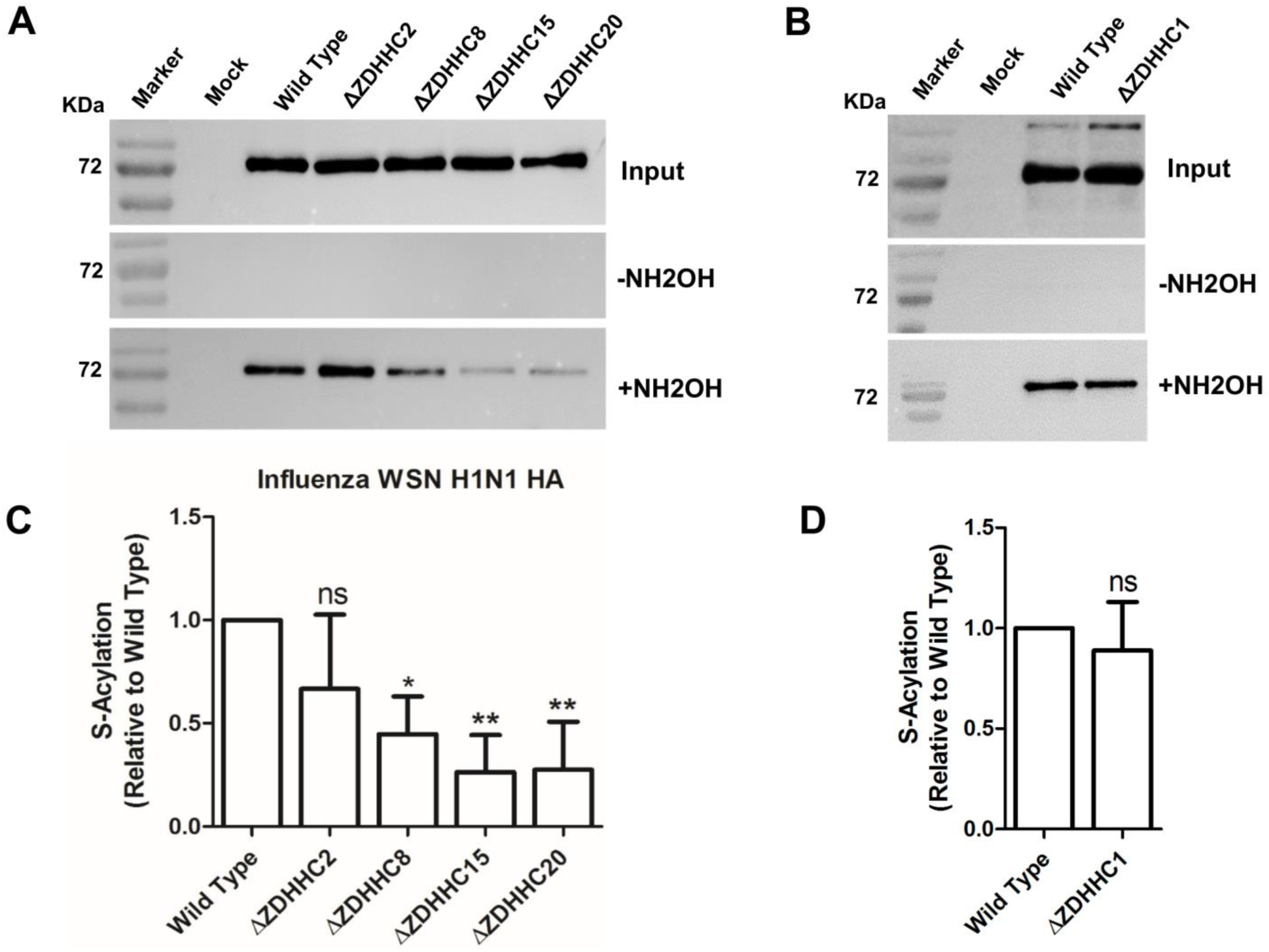
HAP1 cells deficient in ZDHHC2, 8, 15 and 20 show compromised acylation of a group 1 HA. **A+B:** HAP1 cells where the indicated ZDHHCs were knocked-out using the CRISPR/Cas9 technology were infected with the WSN virus, which contains a group 1 HA, at MOI of 1. 24 hours later, acylation of HA was analyzed using Acyl-RAC. **C+D:** Quantification of the result from this and two other identical experiments. Density of the +NH2OH bands was divided by density of bands from the input and normalized to wild type (=1)). The mean ± standard deviation is shown. The asterisks indicate statistically significant differences (*P < 0.05, **P < 0.01) between WT and the deficient cells according. One-way ANOVA followed by Tukey’s multiple comparison test was applied for statistical analysis. ns: not significant.

**Figure 5:**
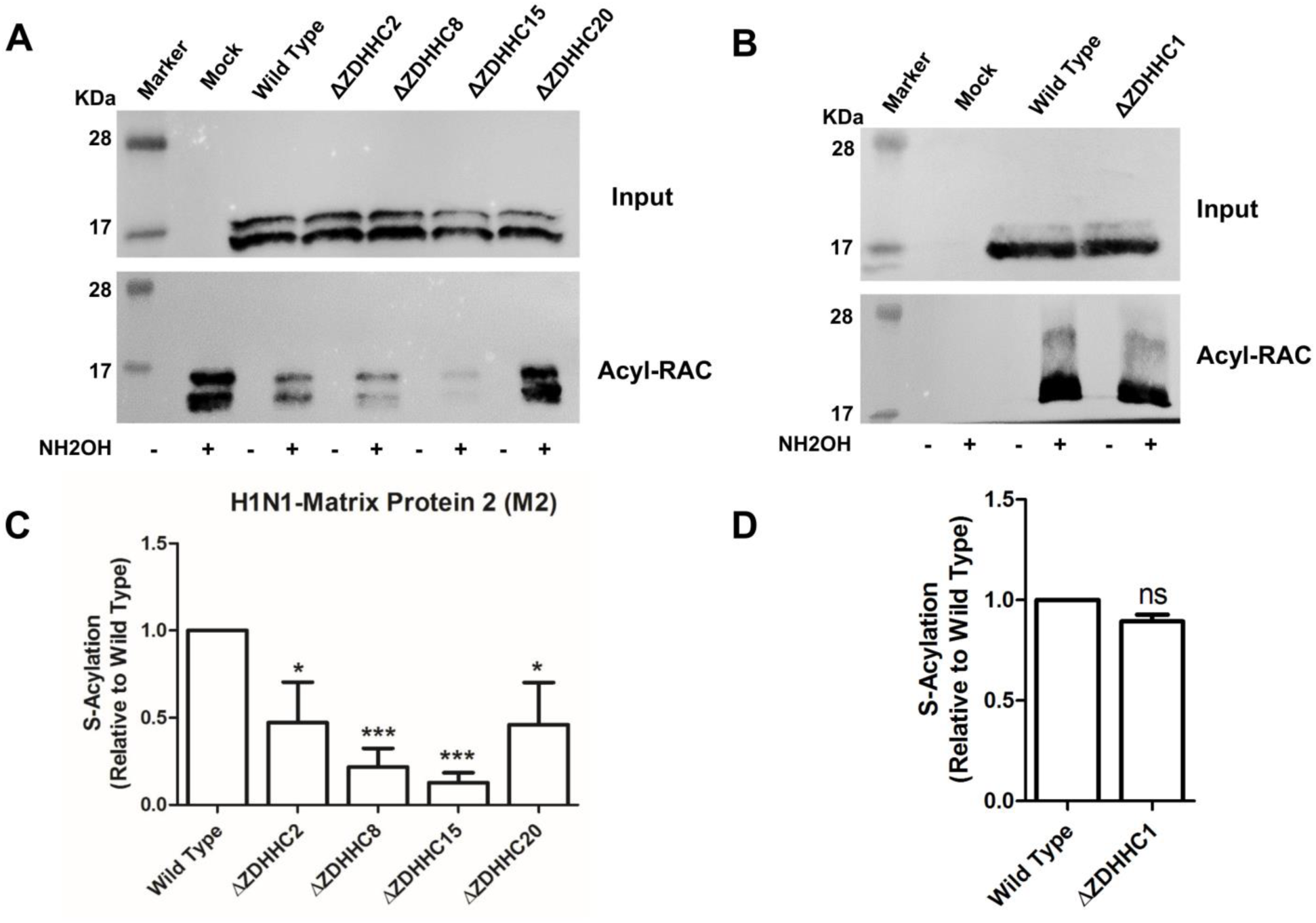
HAP1 cells deficient in ZDHHC2, 8, 15 and 20 show compromised acylation of M2. **A+B:** HAP1 cells where the indicated ZDHHCs were knocked-out using the CRISPR/Cas9 technology were infected with the WSN virus. 24 hours later, acylation of M2 was analyzed using Acyl-RAC and an antibody against M2. **C+D:** Quantification of the result from this and another identical experiment. Density of the +NH2OH bands was divided by density of bands from the input (M2 expression) and normalized to wild-type (=1). The mean ± standard deviation is shown. The asterisks indicate statistically significant differences (*P < 0.05, **P < 0.01, ***P < 0.001) between WT and the deficient cells. One-way ANOVA followed by Tukey’s multiple comparison test was applied for statistical analysis.

**Figure 6:**
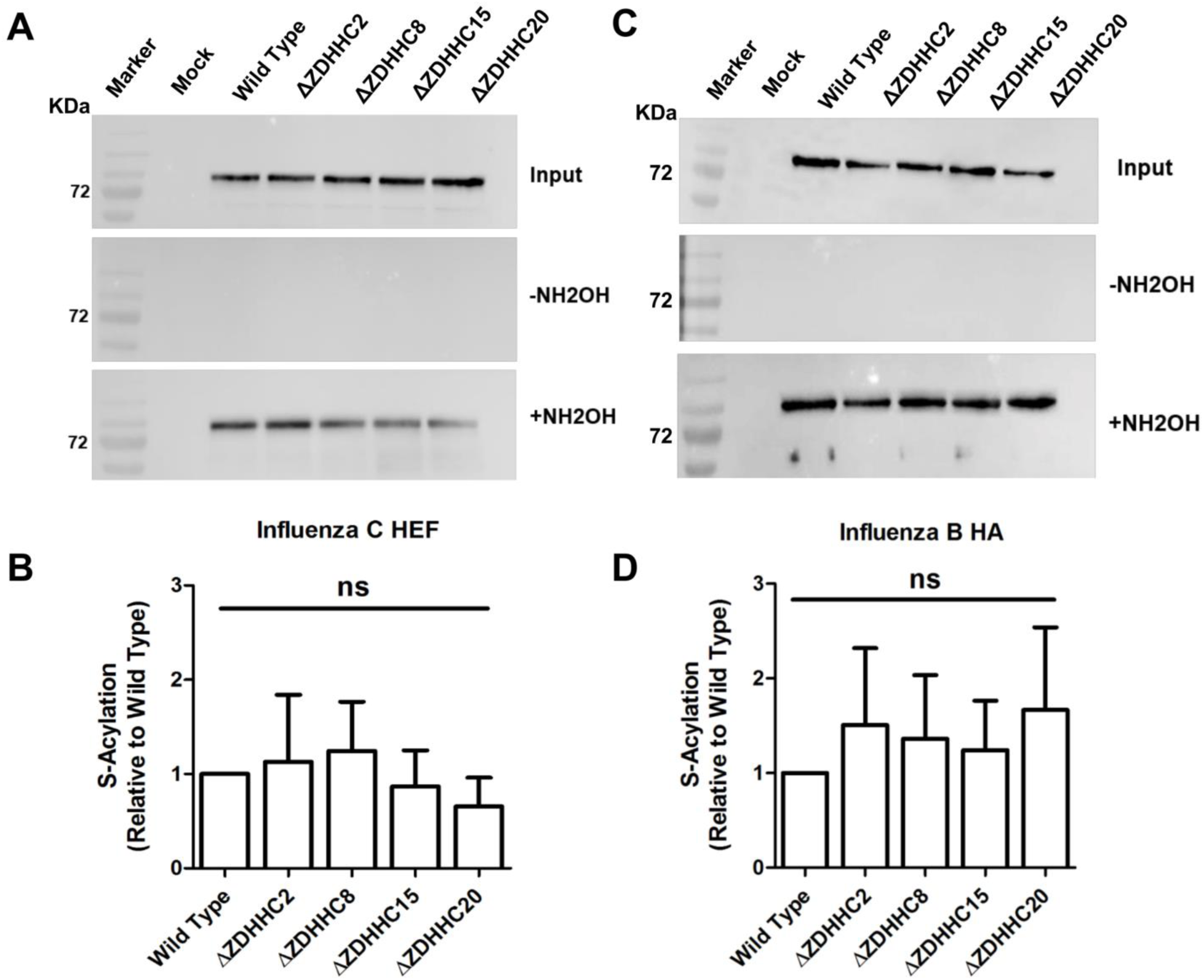
HAP1 cells deficient in ZDHHC2, 8, 15 and 20 exhibit no effect on acylation of HEF from Flu C and HA from Flu B. **A:** HAP1 cells where the indicated ZDHHCs were knocked-out using the CRISPR/Cas9 technology were infected with Influenza C virus. 24 hours later acylation of HEF was analyzed using Acyl-RAC. **B:** Quantification of the result from this and two other identical experiment. Density of bands in the +NH2OH blot was divided by the density of bands from the input (=HEF expression levels) and normalized to wild-type (=1). The mean ± standard deviation is shown. One-way ANOVA followed by Tukey’s multiple comparison test was applied for statistical analysis. **C:** HAP1 cells where the indicated ZDHHCs were knocked out using the CRISPR/Cas technology were infected with Influenza B virus. 24 hours later acylation of HA was analyzed using Acyl-RAC **D:** Quantification of the result from this and two other identical experiments revealed no significant difference.

**Figure 7:**
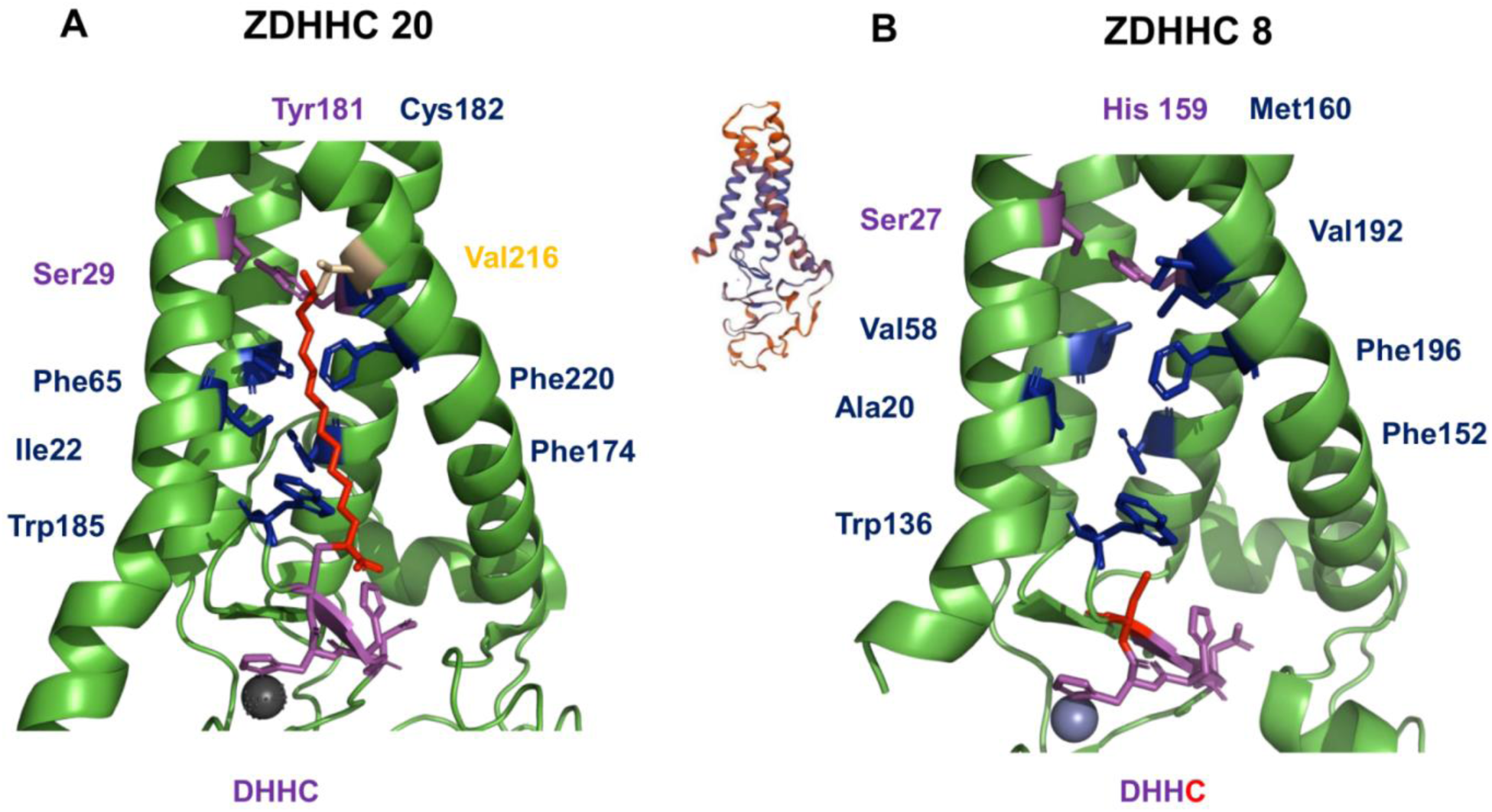
Structure of human ZDHHC20 and a model for ZDHHC8. **A:** The figure shows the structure of the hydrophobic cavity of human ZDHHC20. Ser 29 and Tyr 181 (shown as magenta sticks) form a hydrogen bond that seals the tunnel. Amino acids contacting the acyl chain (red) are shown as blue sticks. They are conserved in ZDHHC2, 15 and 20 except Val216 (orange stick) which is replaced by an alanine in ZDHHC2. The DHHC motif is shown as magenta sticks. The figure was created with PyMOl from pdb-file 6bmm.1.A. **B:** Computational model of ZDHHC8. The structure was created with Swiss-model (https://swissmodel.expasy.org/) using the structure of ZDHHC20 as template. Although the quality of the whole model is quite low, the structure of the transmembrane regions is predicted with higher confidence (blue color in the inset). The amino acids supposed to make contact with the acyl chain are shown as blue sticks. Amino acids Ser 27 and His 159 (magenta sticks) are supposed to close the hydrophobic tunnel.

**Figure 8:**
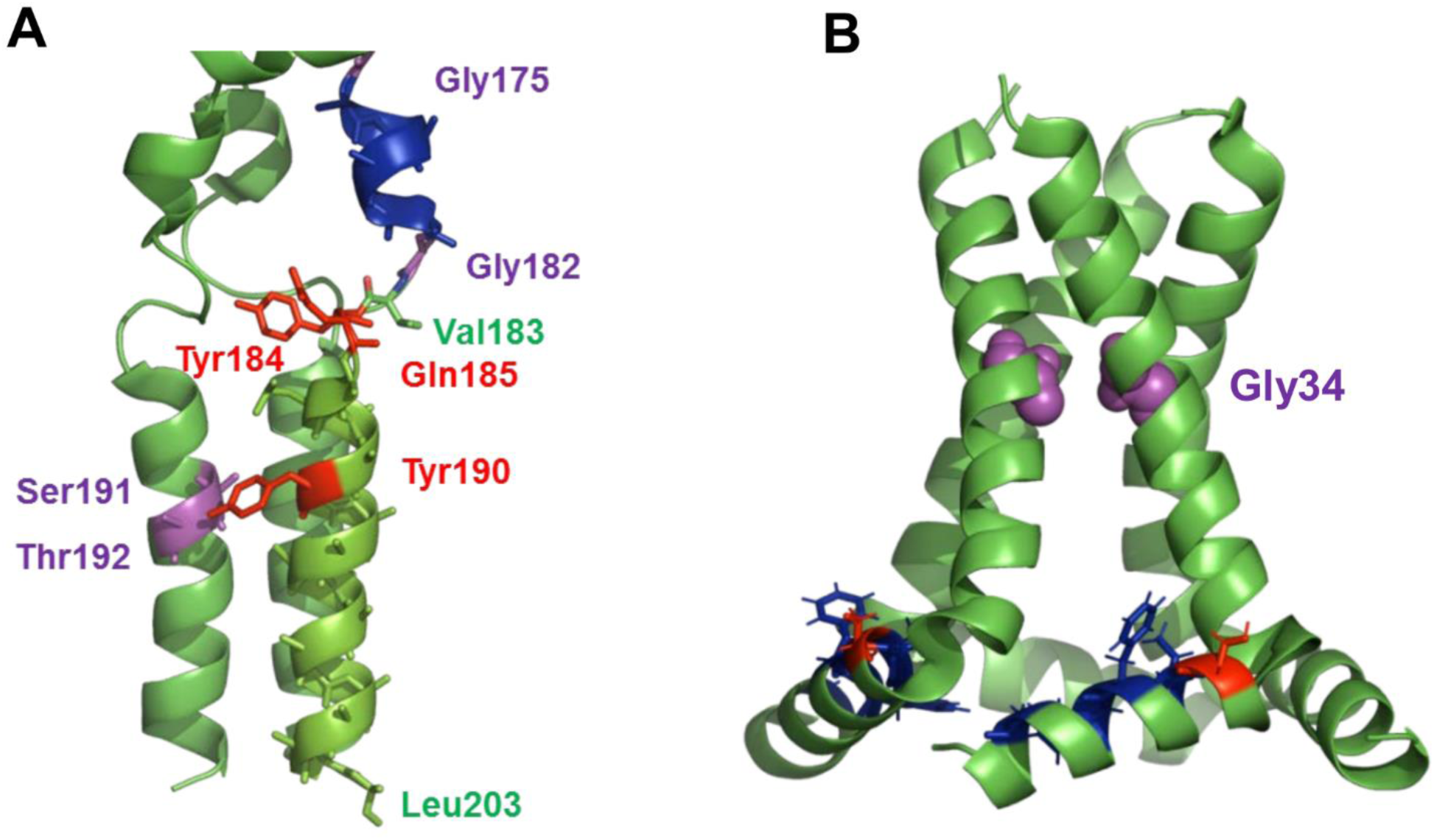
Structure of the viral substrate proteins HA and M2. **A:** C-terminal part of the full-length structure of a group 1 HA. The ectodomain of HA (not shown) is connected by a linker region which contains a short α-helix (blue) to the trimeric transmembrane region. The helix is confined by two glycine residues, which allows the ectodomain to tilt against the membrane anchor. Tyr 190 in the TMR probably forms a hydrogen-bond with Ser 191 or Thr 192 in another helix. The last resolved amino acid in the TMR is Leu 203, the next residue, a glycine causes the helices to splay apart. Note that important residues, such as Tyr190, are not present in group 2 HAs suggesting that it might exhibit a different structure. The figure was created with PyMol from pdb-file 6HJ0. **B:** Structure of amino acids 22-62 of the viral proton channel M2. The tetrameric transmembrane region (residues 26-46) is kinked around the highly conserved glycine 34, which is highlighted as magenta spheres in two TMR regions. Residues 46-49 from a tight turn, which connects the TMR to an amphiphilic helix. Residues interacting with the membrane are shown as blue sticks. Shown as red stick is the acylation site. Cys is replaced by a serine in the peptide used to make the NMR structure. The figure was created with PyMol from pdb-file 2L0J.

## 3. Results

### 3.1 Screening for ZDHHCs involved in acylation of HA using siRNAs

To narrow down the number of palmitoyltransferases that are involved in acylation of HA from Influenza A virus (Flu A) we first performed a broad siRNA screen in HeLa cells. These cells do not produce infectious virus particles [45], but can be transfected with high efficiency. Using real-time PCR, we detected expression of most ZDHHCs (except ZDHHC19), but at different levels (Fig. S1). We transfected Hela cells with a commercial siRNA library against individual human ZDHHCs. This approach is well described and commonly used to identify ZDHHCs involved in acylation of substrates [46]. The same library was already used in HeLa cells to identify ZDHHC6 as the enzyme responsible for palmitoylation of calnexin. It reduces synthesis of individual ZDHHC by ∼70% to 90% [42]. 48 hours later cells were transfected with a plasmid encoding a H7 subtype HA. Cells were lysed and a 10% aliquot was used to compare the expression levels of HA in cells transfected with the different siRNAs using western blotting. The remaining protein sample was equally split and subjected to acyl-RAC (resin-assisted capture) to determine whether acylation of HA is reduced in any of the siRNA transfected cells. Acyl-RAC exploits thiol-reactive resins, which capture SH-groups newly liberated by hyxdroxylamine cleavage of thioester bonds. The other half of the aliquot was treated with Tris-HCL instead of hyxdroxylamine to assess the binding-specificity of the resin. All samples were then subjected to western blotting with an antiserum against the HA2 subunit of H7 subtype HA. Blocking expression of ZDHHC2 and 8 completely inhibited acylation of HA, but not acylation of the endogenous cellular protein caveolin- 1 (Fig. S2). siRNAs against the other ZDHHCs had less pronounced effects. Acylation of HA remained unchanged when using siRNAs against some ZDHHCs, e.g. ZDHHC1, 3, 4; other siRNAs slightly reduced (ZDHHCs12, 14, 19) or enhanced levels of HA’s acylation (ZDHHCs7, 9, 17).

Since the siRNA screen exhibits some technical flaws, such as reduced acylation of HA in the presence of siRNAs against ZDHHC19, although expression of the protein was not detected by qPCR, we aimed to confirm the main findings by metabolic labelling of transfected cells with [^3^H]-palmitate followed by immunoprecipitation of HA and fluorography. Blotting an aliquot of immunoprecipitated HA ensures that similar amounts of the protein are compared in the fluorogram. We used the same siRNAs against ZDHHC2 and 8, but also against ZDHHC15 and 20. Although the latter had no significant effect on acylation of HA in the first screen, they are phylogenetically closely related to ZDHHC2 (Fig. 1a) and therefore might acylate the same substrates. A scrambled siRNA and a siRNA against ZDHHC1, which had no effect in the siRNA screen, were used as negative controls. Results shown in Fig.1b revealed that [^3^H]-palmitate labelling of HA is strongly inhibited by siRNAs against ZDHHC2, 15 and 20 and clearly reduced by siRNAs against ZDHHC8.

### 3.2 Knockout of ZDHHC2, 8, 15 or 20 in HAP1 cells reduces acylation of HA of influenza A virus

We next used commercial HAP1 cells, where ZDHHCs2, 8, 15 and 20 were knocked out using the CRISPR/Cas9 technology. HAP1 cells are a human cell line derived from the chronic myelogenous leukemia cell line KBM-7, which have a single copy of almost every chromosome and are therefore ideally suited for the knockout technology. Wild-type HAP1 cells express a similar set of ZDHHCs as HeLa cells, only the expression levels vary between cell types (Fig. S1). Next, we compared the mRNA levels between the CRISPR/Cas9 treated and wild type HAP1 cells. Whereas expression of the corresponding mRNAs is decreased in ΔZDHHC2 and ΔZDHHC20 to 50 and 20%, respectively, ΔZDHHC8 and ΔZDHHC15 cells synthesize slightly more of the relevant mRNAs (Fig. s3). Since the ΔZDHHC HAP1 cells were generated using single guide RNA intended to cause only small deletions or insertions (see Fig. s4 for details), it is not unexpected that mRNAs are still transcribed.

Therefore, we analysed by sequencing of chromosomal PCR products whether the ZDHHCs contain frame-shift mutations within their coding exon. All sequencing chromatograms (Fig. S4) clearly show a difference in the nucleotide sequences within the Cas9 binding site between wild type and ZDHHC knockout cells. Since the nucleotide differences are all located either before or within the CRD-DHHC domain it is unlikely that the knockout cells are able to synthesize a functional ZDHHC protein, even if the mRNAs are translated.

HAP1 cells can be infected with Influenza virus and synthesize HA, which is transported to the plasma membrane as demonstrated by binding of erythrocytes to infected cells. However, HAP1 cells do not release virus particles as analysed by hemagglutination and plaque assays (not shown). Acyl-RAC used on cells infected with the virus that expresses the HA used in the siRNA screen (A/chicken/Rostock/8/1934 (H7N1)) confirmed the results. Acylation of HA is strongly inhibited in ΔZDHHC2, ΔZDHHC15 and ΔZDHHC20 and reduced in ΔZDHHC8 knockout cells, whereas very little (if any) effect was observed in ΔZDHHC1 cells, which were used as control (Fig. 2a-b). Quantifying relative acylation of HA (band densities of the acylation signal relative to total expression) from this and two identical experiments revealed that acylation is reduced by ∼75% in ΔZDHHC15 and ΔZDHHC20 and to ∼65% in ΔZDHHC2 cells. ΔZDHHC8 cells show the tendency to less efficiently acylate HA, but the mean (80%) was not statistically significant different from wild type cells (Fig. 2c). Since ZDHHC8 is phylogenetically related to ZDHHC5, we also tested ΔZDHHC5 cells, but no reduction in acylation of HA was observed (Fig S5).

### 3.3 Co-localization of ZDHHC2, 8, 15 and 20 with HA in a human lung cell line

We investigated next whether the ZDHHCs we identified to be involved in acylation of HA are expressed in a cell line relevant for replication of Influenza virus, namely human lung epithelial A549 cells. Real-time PCR revealed that all ZDHHC (except 19) are expressed (Fig. s1). The relative abundance of the relevant transcripts is the same in each cell type we used in this study. From the three phylogenetic related ZDHHCs2, 15 and 20 exhibits ZDHHC15 the lowest and ZDHHC2 the highest expression level in each cell. ZDHHC8 is expressed at a similar level as ZDHHC2 (Fig. s1).

We then asked whether the intracellular localization of the identified ZDHHCs is consistent with the intracellular site of HA’s acylation. Since HA pulse-labeled with [^3^H]-palmitate is already trimerized, but its carbohydrates are still Endo-H sensitive it can be concluded that acylation occurs in the late-ER, in the ERGIC and/or in the cis-Golgi region [47]. ZDHHC2, 8 and 15 were previously localized in transfected HEK293T cells to the cis-Golgi, whereas ZDHHC2 also co-localizes with an ER marker [33]. In the same study, ZDHHC20 was detected at the plasma membrane, but others reported that it is also present in a perinuclear compartment [33, 48]. To investigate whether HA co-localizes with the ZDHHCs we co-transfected A549 cells with the H7 subtype HA and with either ZDHHC2, 8, 15 and 20, each fused to a C-terminal HA-tag [41]. Staining of permeabilized cells with an anti-HA-tag antibody showed an intracellular, reticular staining pattern for each ZDHHC, often more concentrated near the nucleus and exhibiting brighter spots throughout the cell (Fig. 3). Staining of HA with anti-Influenza virus serum revealed that it co-localizes with each ZDHHC protein, but only inside cells, not at the plasma membrane where HA is also abundant. In neurons ZDHHC2 cycles between the plasma membrane and endosomes [49]. However, since HA is not endocytosed in transfected cells [50], it is unlikely that the intracellular co-localization of ZDHHC2 with HA takes place in endocytic vesicles. Quantification of the degree of co- localization of the two fluorophores using the Pearson’s correlation coefficient revealed that 60% to 75% of HA molecules co-localize with individual ZDHHCs (see legend of figure 3 for details). However, this is not necessarily due to long-lasting binding of HA to individual ZDHHCs, but rather indicates that the ZDHHCs are present at membranes (or membrane domains) of the exocytotic pathway, the site where acylation of HA occurs [47].

### 3.4 HAs of both phylogenetic groups and the viral proton channel M2 are acylated by a similar set of ZDHHC proteins

HA proteins from the various Influenza strains are divided into two phylogenetic groups; the H7 subtype HA from an avian virus we investigated so far belongs to group 2. For a group 1 HA a 3D-structure of most parts of the transmembrane region was resolved by Cryo-EM [36]. However, critical amino acids are not conserved in group 2 HAs (see Fig. 8) suggesting that group 1 and 2 HAs might exhibit different structures. We therefore asked whether they are acylated by the same set of ZDHHC proteins. We infected the ΔZDHHC HAP1 cells with WSN virus (A/WSN/33, H1N1), which was originally isolated from a human patient and performed Acyl-RAC experiments. Quantification of the results from three virus infections revealed that knockout of ZDHHC15 and 20 had the strongest effect (80% reduction) on acylation of HA. Abolishment of the ZDHHC8 protein also led to a significant reduction in HA’s acylation to 30%. ΔZDHHC2 cells showed the tendency to less efficiently acylate HA, but since there were large variation between experiments the mean (50%) was not statistically significant different from wild type cells (Fig. 4). ΔZDHHC1 and ΔZDHHC5 cells had very little (if any) effect on acylation of the group 1 HA (Fig 5b+d and Fig. S5)

Next, we asked whether M2, the second palmitoylated protein of Influenza virus is also acylated by the same set of ZDHHC proteins. Acyl-RAC on HAP1 cells infected with the WSN virus revealed that acylation of M2 was affected in each ZDHHC knockout cells, but to a different extent compared to acylation of HA (Fig. 5). M2’s acylation is affected the most in ΔZDHHC8 and ΔZDHHC15 cells, less prominently in ΔZDHHC2 and ΔZDHHC20 cells. The reduction in acylation of M2 was statistically significant for each ΔZDHHC knockout cell. Again, ΔZDHHC1 and ΔZDHHC5 cells had no effect on acylation of M2 (Fig 5b+d and Fig. S5). In summary, the acylated proteins of Influenza A virus are modified by the same set of ZDHHC proteins, but some variation in the relevance of each ZDHHC for acylation of group 1 and 2 HAs and M2 apparently exist.

### 3.5 ZDHHC 2, 8, 15 and 20 have no effect on acylation of HA of Flu B and HEF of Flu C

HA of Influenza B virus and HEF of Influenza C virus exhibit the same overall three*-*dimensional folding of their polypeptide chains and play identical roles during virus replication as HA of Flu A, but they differ in their acylation pattern. We were therefore interested whether all hemagglutinating glycoproteins of Influenza viruses are acylated by the same ZDHHC proteins. Only ΔZDHHC15 and ΔZDHHC20 cells infected with Influenza C virus revealed a small (but not statistically significant) reduction in acylation of HEF, whereas ΔZDHHC2 and ΔZDHHC8 HAP1 cells exhibit no effect (Fig. 6a+b). This result is not unexpected since none of the ZDHHCs identified so far exhibits a very high specificity for stearate (see discussion) and we might have missed a stearate-specific ZDHHC in our siRNA screen since it reduces total acylation of HA at most by one-third. Surprisingly, however, knockout of ZDHHC2, 8, 15 and 20 did also not reduce acylation of HA in HAP1 cells infected with Influenza B virus. The mean of three experiments showed that acylation of HA was even slightly (but not statistically significant) enhanced (Fig. 6c+d).

Thus, HA of Flu B and HEF of Flu C are apparently acylated by other, yet to be identified ZDHHC proteins than HA of Flu A. These results also clearly demonstrate that blocking the expression of ZDHHC2, 8, 15 and 20 in HAP1 cells does not cause a general reduction of acylation of transmembrane proteins that are transported along the exocytic pathway and thus the effect we observe on HA and M2 of Influenza A virus is specific.

## 4. Discussion

### 4.1 ZDHHC2, 8, 15 and 20 are involved in acylation of HA and M2 of influenza A virus

Here we show that expression of ZDHHC2, 8, 15 and 20 is required for acylation of HA and M2 of Influenza A virus. This was demonstrated using a siRNA approach in influenza HA-transfected HeLa cells (Fig.1) and confirmed with CRISPR/Cas9 technology in virus-infected HAP1 cells (Fig. 2). Reduction in acylation of HA was analyzed both with [^3^H]-palmitate-labelling and Acyl-RAC assays. The differing result for siRNAs against ZDHHC15 and 20 between the Acyl-RAC (Fig. S1) and the [^3^H]- palmitate assays (Fig. 1B) might be due to the fact that different populations of HA molecules are detected. Acyl-RAC determines all HA molecules synthesized after transfection, i.e. in a time period of 24 hours. In contrast, [^3^H]-palmitate detects only HA molecules synthesized in a time span of two hours and 24 hours after transfection. Thus, one might speculate that ZDHHC15 and 20 preferentially acylate HA molecules synthesized late after transfection, for example because their activity might be upregulated or because the capacity of ZDHHC2 and 8 might be saturated at late time points.HA co-localizes with ZDHHC2, 8, 15 and 20 at membranes of the exocytic pathway in A549 cells (Fig. 3), a human epithelial lung cell line, which is consistent with the intracellular site of acylation of HA. Moreover, since quantitative PCR (qPCR) revealed that ZDHHC2, 8, 15 and 20 (as well as every other ZDHHC except ZDHHC 19) are expressed in A549 cells (Fig. S1) the ZDHHCs we identified are probably also relevant for acylation of HA in human lung cells. Unfortunately, since HeLa and HAP1 cells do not produce infectious virus and A549 or MDCK cells, which support virus replication, are hard to transfect we could not use siRNA screens to test whether knockdown of these ZDHHCs reduces virus titers as the essential nature of the protein modification would suggest.

Based on co-expression experiments of various types of substrate proteins with mammalian ZDHHCs in yeast cells it was proposed that ZDHHC2 and ZDHHC20 (but not ZDHHC15) have high activity towards integral membrane proteins [51]. However, ZDHHC2, 8, 15 and 20 are not involved in acylation of HA of Flu B and HEF of Flu C, although all hemagglutinating glycoproteins of Influenza viruses are typical type 1 transmembrane proteins, which exhibit a similar 3D-structure. Furthermore, each of them is transported along the exocytic pathway to the plasma membrane and thus passes the same set of ZDHHC proteins.

One might speculate that the different ZDHHCs hijacked by Influenza A and Influenza B and C viruses might be the result of their different host origins. The reservoirs of Influenza B and C viruses are almost exclusively humans. In contrast, all types of Influenza A viruses constantly circulate in water birds, from where some of them occasionally spread to other animals, such as poultry and swine or to humans. Thus, it might be that the viruses adapted independently from each other to a different set of ZDHHC proteins, i. e. Influenza A viruses adapted in birds to ZDHHC2, 8, 15 and 20, Influenza B and C viruses in humans to other unknown ZDHHCs. This seems contradictory to the finding that the human WSN virus (H1N1) uses the same set of ZDHHCs as the avian fowl plaque virus (H7N1). However, WSN is a successor of the virus that caused the 1918 pandemic, which is likely the result of direct transmission of an avian virus into the human population [52]. The avian viruses likely use the same ZDHHC proteins in both avian and mammalian cells, at least HA from WSN virus purified from embryonated hen’s eggs and from mammalian cells are both stoichiometrically acylated. Minor differences were identified in HA’s fatty acid pattern; more stearate was attached if the virus was grown in mammalian (20%) compared to avian cells (10%) [12].

We do not want to exclude that other ZDHHCs might also contribute to acylation of HA since the initial screen revealed reduction in HA’s acylation by siRNAs directed against various ZDHHC proteins. From an evolutionary point of view, it would be unfavorable for a virus to rely completely on just one (or a few) ZDHHC protein(s) to catalyze a protein modification that is essential for its replication. Such a specialization might restrict virus tropisms to cells and organisms where this ZDHHC is highly expressed and functionally active since a large number of HA molecules need to be acylated. One virus particle contains ∼500 trimeric HA spikes and 50 M2 proton channels [53], which correspond to 200 M2-linked and 4500 HA-linked fatty acids because every acylation site is completely filled [10]. Since each cell releases up to 5000 particles in ∼10 hours 20-30 million fatty acid bonds need to be catalyzed in one viral replication cycle. This estimation does not even take into account the large number of HA and M2 molecules still present inside dead cells, which are not able to release more particles. We thus assume that HA has adapted to a ZDHHC machinery that has a high capacity.

The reduction of acylation of HA by eliminating a single ZDHHC is at least 50% and thus stronger than expected if the four ZDHHCs are redundant. We do not want to conclude that each of the identified ZDHHCs actually transfers fatty acids to HA. It is possible that they are part of a palmitoylation cascade, similar to the one involved in acylation of ER-resident protein folding factors [54]. In that case, ZDHHC6 catalyzes fatty acid transfer to the protein substrate, but its activity is regulated by ZDHHC16-catalyzed palmitoylation of three cytoplasmic cysteines in ZDHHC6. This generates three versions of the enzyme: non-palmitoylated ZDHHC6 is inactive, palmitoylation at the first site creates a highly active ZDHHC version that is rapidly degraded and acylation at at least one of the two other cysteines generates a moderately active and stable version. Using mass spectrometry both ZDHHC8 and ZDHHC20 appeared to be S-acylated on three cysteine residues within a CCX7–13C(S/T) motif in their cytoplasmic tails suggesting that they might be regulated by acylation [55, 56]. However, it seems unlikely that all four identified ZDHHCs are part of a palmitoylation cascade. One might thus speculate that some of them work synergistically with each other, i. e. acylation of a single cysteine in HA by a certain ZDHHCs makes another cysteines accessible to another ZDHHC enzyme. More sensitive methods to profile the acylation status of HA (including the type of attached fatty acid) are required to verify these assumptions, but our MS approach is currently feasible only for HA present in virus particles, not for intracellular HA [12].

### 4.2 Fatty acid specificities of ZDHHCs involved in acylation of HA

Do the hitherto determined lipid specificities of the ZDHHCs involved in acylation of HA and M2 correspond to the peculiar fatty acid pattern determined for these proteins? ZDHHC2, 15 and 20 use myristate, palmitate and stearate in the auto-acylation reaction and transfer it to protein substrates, but with different efficiency. ZDHHC2 does not show a significant preference for any of the acyl chains, whereas ZDHHC15 and 20 prefer the shorter acyl chains, even myristate is preferred over palmitate [38, 39]. However, we never detected myristate as fatty acid bound to HA or to any other viral glycoprotein [10, 57].

One factor limiting transfer of myristate to viral proteins might be a low Myr-CoA concentration inside cells. This might be assumed because myristate is a rare acyl-chain, both as free fatty acid and bound to phospholipids. However, at least in HEK293 cells the Myr-CoA content is almost half of the Pal-CoA content and only slightly lower than the amount of Stear-CoA [38]. Nonetheless, the local concentration of Myr-CoA in the vicinity of a ZDHHC protein might be lower and thus limiting myristate transfer to proteins. Pal-CoA has been shown to insert into artificial membranes [58], but Myr-CoA, due to its shorter acyl-chain might have a lower propensity to interact with bilayers, which is reminiscent of myristoylated peptides that have a much lower membrane binding affinity compared to palmitoylated peptides [59]. Alternatively, acyl-CoA binding proteins (ACBP), ubiquitous, mostly cytosolic polypeptides, might control the availability of acyl-CoAs for ZDHHCs. They occur as single domain polypeptide, but the ACBP domain is also present in ∼50 different proteins. They bind acyl-CoAs with high affinity (K_D_ 1-10nM) which keeps the intracellular concentration of free acyl-CoAs in the low nM range. Most of them bind a variety of long chain acyl-CoAs, but at least the parasite *Plasmodium falciparum* encodes an ACBP that is specific for Myr-CoA [60]. It is tempting to speculate that ACBPs might sequester Myr-CoA, which is then not available for ZDHHC proteins. Alternatively, ACBPs might play an active role, e.g. transfer specific lipid substrates from the cytosol to certain ZDHHCs.

The crystal structure of human ZDHHC20 provided a molecular explanation which fatty acids are accepted in the acylated enzyme intermediate [39]. The acyl chain is inserted into a hydrophobic cavity formed by all four transmembrane regions. ZDHHC20 contains at the narrow end of the cavity the small amino acid Ser at position 29 and the large amino acid Tyr at position 181 that form a hydrogen bond which effectively closes the groove (Fig. 7a). Most ZDHHCs contain either two bulky residues, one bulky and one small or two small amino acids at the homologues position and it was postulated that the presence of two large residues would limit the use of stearate. Indeed, exchange of Ser 29 by Phe reduced and of Tyr 181 by Ala enhanced the usage of stearate by ZDHHC20. A similar observation was also reported for the ZDHHC3 enzyme [38].

Based on this model we asked whether the other ZDHHCs we identified to be involved in acylation of HA also accept stearate as lipid donor. Sequence comparison shows that ZDHHC2 and 15 also contain a serine and a tyrosine at the end of the cavity and all other (except one) hydrophobic residues lining the cavity and contacting the acyl chain are conserved between ZDHHC2, 15 and 20. Only ZDHHC2 contains an alanine instead of a valine at position 216, which is located near the end of the tunnel. Since alanine has a shorter side chain, this might explain the increased usage of stearate in autoacylation of ZDHHC2 relative to ZDHHC15 [38].

Lipid preferences have not been experimentally tested for ZDHHC8. We therefore created a 3D-model of ZDHHC8 using the crystal structure of ZDHHC20 as template (Fig. 7b). It revealed identical or similar amino acids in most parts of the hydrophobic cavity of both proteins: Trp136 at the entrance of the tunnel, Phe152, Val192 and Phe196 in TMR 3 and 4 and hydrophobic, but shorter amino acids (Val20, Ala58) in TMR 1 and 2. A cysteine at the end of the cavity is exchanged to methionine 160 in ZDHHC8. Interestingly, the cavity is sealed by two small amino acids, Ser27 and His159 and thus one might speculate that ZDHHC8 might exhibit a higher preference for stearate than ZDHHC20.

In summary, the ZDHHCs we identified to be involved in acylation of HA of Flu A are apparently able to transfer both palmitate and stearate to a substrate. Thus, site-specific acylation of HA can currently not be explained by the activities of two different, acyl-chain specific enzymes. However, it seems possible that the ZDHHC specific for stearate has not been identified in our screen and this ZDHHC might cause the residual acylation of HA in the ΔZDHHC2, 8, 15 and 20 knockout cells. Unfortunately, the mass spectrometry method we used to precisely determine the acyl chain pattern of HA in virus particles [10] can currently not be applied to intracellular HA due to limited sample amounts. However, assuming that the current model for acyl-chain selectivity applies to all ZDHHC proteins, it is hard to envision a ZDHHC that transfers only stearate. Lengthening the tunnel might allow better access of stearate to the hydrophobic cavity but does not discriminate against the shorter palmitate chain. Interestingly, HEF expressed in insect cells as well as the total pool of S-acylated proteins from these cells contain only minor amounts of stearate [61] suggesting that ZDHHCs have evolved to higher complexity in mammalian cells.

### 4.3 Protein substrate recognition of ZDHHCs involved in acylation of HA and M2

It was recently proposed that acylation of transmembrane proteins occurs whenever a cysteine is accessible, i.e. located maximally 5-6 helix residues into the inner membrane leaflet [62]. This is consistent with the observation that cysteines located in the middle of the transmembrane region of certain HA-subtypes are not acylated [9]. Since even prokaryotic proteins with cysteines inserted near the transmembrane region become palmitoylated when expressed in mammalian cells it was also proposed that acylation is a random (stochastic) process that does not depend on recognition of a specific sequence or structural motif [62]. However, at least HA of Influenza A viruses must exhibit a certain feature recognized by ZDHHC2, 8, 15 and 20, which is absent in HA of Influenza B virus.

Since the largest part of HA is exposed to the lumen of the ER/Golgi and hence not accessible for a ZDHHC, only a short linker (8 residues), the transmembrane region (26-30 residues) and the small cytoplasmic tail (10-11 amino acids) might contain such a signal (table 1). Common to the cytoplasmic tails of both HAs are two conserved hydrophobic amino acids surrounding the palmitoylated cysteine at the C-terminus (ICI in Flu A, ICL in Flu B). This hydrophobic patch might cause the cytoplasmic tail to run parallel to the membrane bilayer. Exchange of one hydrophobic by a hydrophilic amino acid prevents virus replication, but not palmitoylation of expressed HA from Flu A [12]. Also, otherwise, the HA tails of Flu A and Flu B contain similar amino acids surrounding the acylation sites, such as asparagine in addition to positively and negatively charged residues. Unique for HA of Flu A is a completely conserved glycine in the cytoplasmic tail, which, however, can be exchanged without affecting the stoichiometry of acylation of HA. Since exchange of other amino acids in the cytoplasmic tail also had no effect on acylation and little influence on virus replication [12, 23], it seems likely that putative acylation signals are rather located in the transmembrane region.

**Table 1:** Amino acids near the acylation sites of viral substrate proteins. Amino acid sequences of the linker, transmembrane region (underlined) and cytoplasmic tail of the viral proteins analyzed in this study. Cysteines acylated with palmitate and stearate are highlighted in yellow and green, respectively. Glycines in the transmembrane region of HA and M2 of Flu A and HEF of Flu C are indicated as red letters. The C-termini of HAs contain conserved hydrophobic residues around the last palmitoylated cysteine (highlighted in grey). This hydrophobic patch might cause the cytoplasmic tail to run parallel to the membrane.

|  |  |
| --- | --- |
| **Flu A HA/1** | GVKLESMG <u>VYQILAIYSTVASSLVLLVSL</u><u>GAISFWM</u><u>CS</u> NGSLQ<u>CR</u><u>CI</u> |
| **Flu A HA/2** | PVKLSSGYKD <u>IILWFSF</u><u>GASCFLLLAIAM</u><u>GLVFI</u><u>CV</u> KNGNMR<u>CT</u><u>CI</u> |
| **Flu B HA** | SLNDDGLD <u>NHTILLYYSTAASSLAVTLMIAIFIVYMVS</u> RDNVS<u>CS</u><u>ICL</u> |
| **Flu C HEF** | DTKIDLQSD <u>PFYW</u><u>GSSL</u><u>GLAITATISLAALVIS</u><u>GIAI</u><u>C</u> RTK |
| **Flu A M2** | NDSSDP <u>LVIAANII</u><u>GILHLILWIL</u> DRL FFK<u>C</u>IYRRLKYGLKR.. |

The structure of the outer part of the TMR has been resolved by Cryo-EM for a group 1 HA from Influenza A virus (Fig 8a). A flexible linker that contains a five-residue α-helix connects the ectodomain to the TMR, which forms a bundle of three α-helices. A leucine is the last residue present in the resolved structure. The next residue is a glycine which causes the chains to splay apart and become less ordered, suggesting that the glycine causes a kink in the TMR α-helix [36]. No such detailed structures are available for any other HA, but most likely, they also form trimeric helices [63]. Interestingly, HA from every subtype of Flu A contains a glycine in the middle of the TMR, whereas none of the Flu B HAs present in the database contain any glycine in their TMR (table 1). In our previous study, we were not able to change this glycine in HA of Flu A to a large and hydrophobic isoleucine, since the nucleotides rapidly reverted to a triplet encoding a serine, an amino acid also having a short side chain. The resulting viruses contain stoichiometrically acylated HA, but with marginally increased stearate content [12, 23].

Does M2, the other identified substrate of ZDHHC2, 8, 15 and 20, exhibit a similar feature? The NMR structure of the tetrameric proton channel embedded in a lipid bilayer revealed that the acylation site is located at the beginning of a 15 amino acid long amphiphilic helix that runs perpendicular to the membrane. The helix is connected by a short and tight linker to an 18 residue long transmembrane region, which is indeed kinked around a conserved glycine (Fig. 8b, [37]). We thus suggest that ZDHHCs involved in acylation of HA and M2 of Influenza A virus might recognize proteins with kinked transmembrane regions. This might allow the protruding part of the TMR of a prospective substrate protein to interact with specific ZDHHCs, perhaps by insertion into a pocket between two of their transmembrane regions. Alternatively, due to the absence of a side chain, a glycine in the TMR helix of a substrate provides a flat surface for tight packing of a large, hydrophobic side chain present in the TMR of certain ZDHHC proteins. Other transmembrane proteins which are acylated by ZDHHC2 (tetraspanins CD9 and CD151, [64]) or by ZDHHC20 (interferon induced transmembrane protein 3, [65]) at membrane-near cysteine also contain glycine residues in at least one of their transmembrane regions. However, since ZDHHC2, 8, 15 and 20 do not only acylate transmembrane, but also peripheral membrane proteins there must be other molecular features that these ZDHHCs recognize [27].

## Abbreviations

DHHC-protein: polytopic membrane proteins containing an Asp-His-His-Cys motif;
ER: endoplasmic reticulum;
HA: hemagglutinin;
TMR: transmembrane region.

## Author Contribution

M. V. and G. G. designed the study, M. G. and L.A. performed the experiments. M. G. and M. V. wrote and L. A. and G. G. edited the manuscript.

## Acknowledgements

We thank Ralf Wagner and Hans-Dieter Klenk (Virology, Marburg) for providing the H7 subtype HA, Florian Krammer (Mount Sinai School of Medicine, New York) for the antibodies against HA of Influenza B virus and Masaki Fukata (National Institutes for Physiological Sciences, Japan) for HA- tagged mouse ZDHHC clones.

## Supplementary Figures and Tables

**Figure S1:**
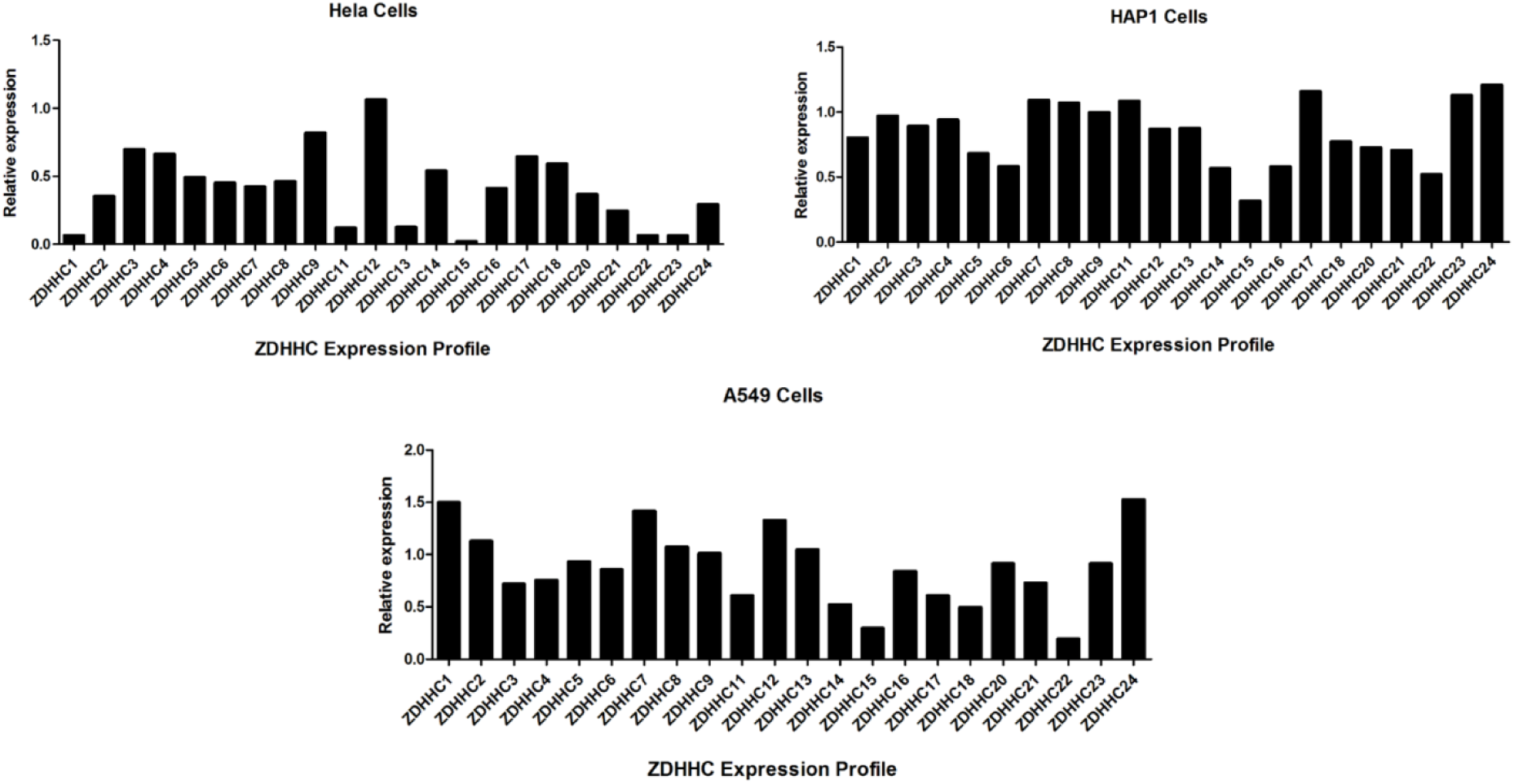
Expression profile of ZDHHC enyzme family members by quantitative real-time PCR (qPCR) for the cell lines used in this study. mRNA levels of HeLa and HAP1 cells were normalized using human TATA-binding protein and β- glucuronidase while expression in A549 cells was normalized to human GAPDH. All ZDHHCs (except ZDHHC19) are expressed in HeLa, HAP1 and A549 cells.

**Figure S2:**
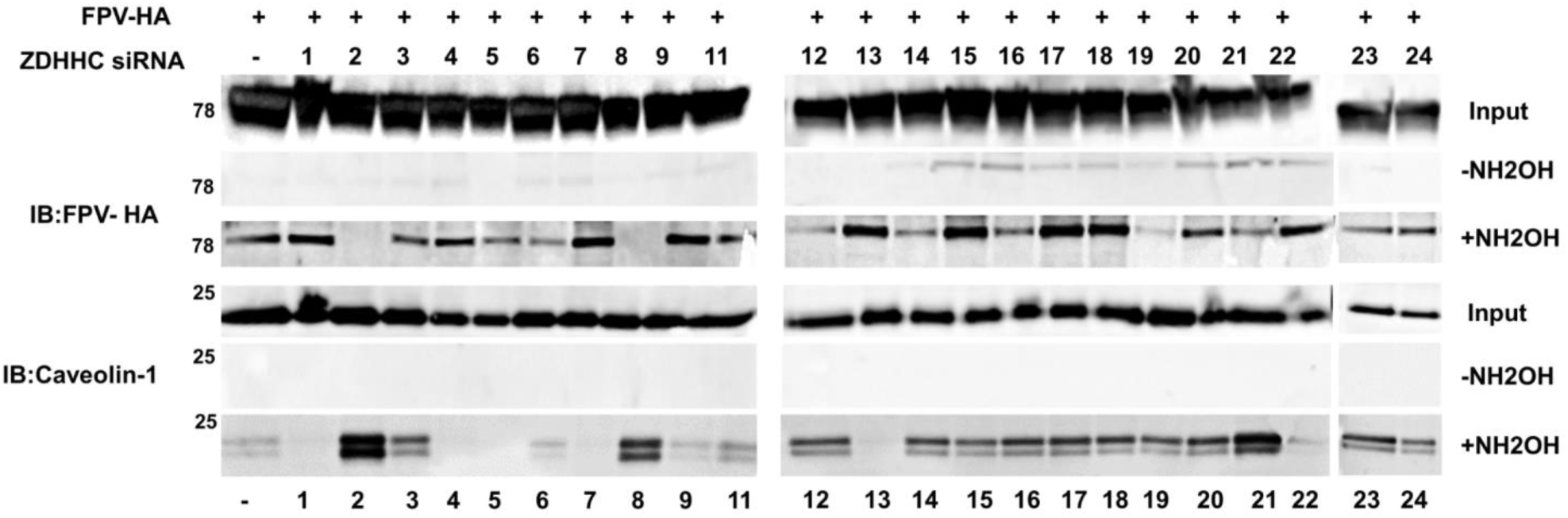
siRNAs against ZDHHC2 and 8 block acylation of HA in transfected HeLa cells. HeLa cells were transfected with siRNAs against all human ZDHHCs and 48 hours later transfected with a plasmid encoding H7 subtype HA from a variant of FPV. After 24 hours, cells were lysed and subjected to acyl-RAC and immune-blotting (IB). First with antiserum against the HA2 subunit (upper panel), then the membrane was again blotted with antibody against caveolin-1 as cellular control (lower panel). NH2OH: samples treated (+) or not treated (-) with hydroxylamine to cleave thioester-bound fatty acids. Input: western blot of 10% of the lysate to check expression levels of HA. 78 and 25 indicates the mobility of the molecular weight marker. The blots on the left were exposed for the same time as the blots in the middle and on the right.

**Figure S3:**
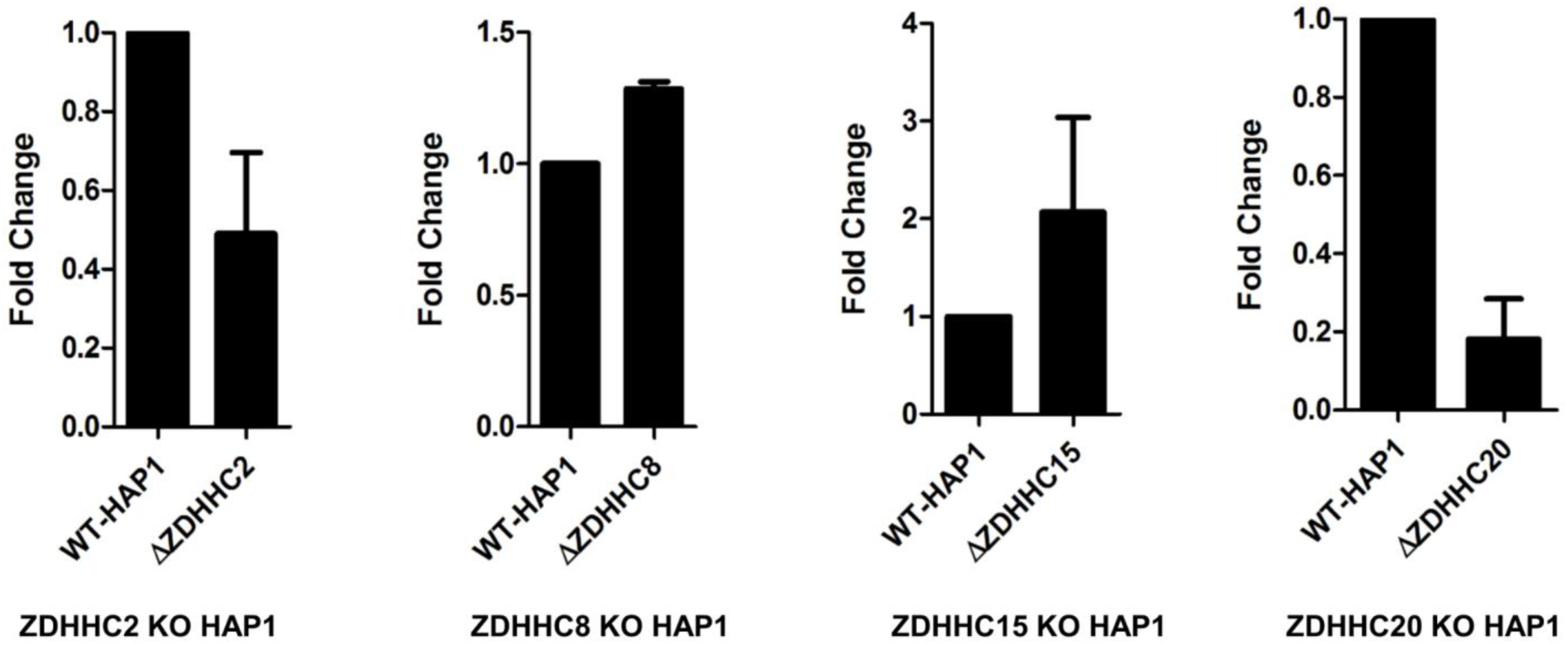
Expression of relevant ZDHHCs in CRISPR/Cas9 knockout relative to wild type HAP1 cells. Graphs showing relative expression level of indicated ZDHHCs determined with qPCR for the specific ZDHHC transcripts, normalized to GAPDH housekeeping gene. Data are the average of two independent experiments.

**Figure S4:**
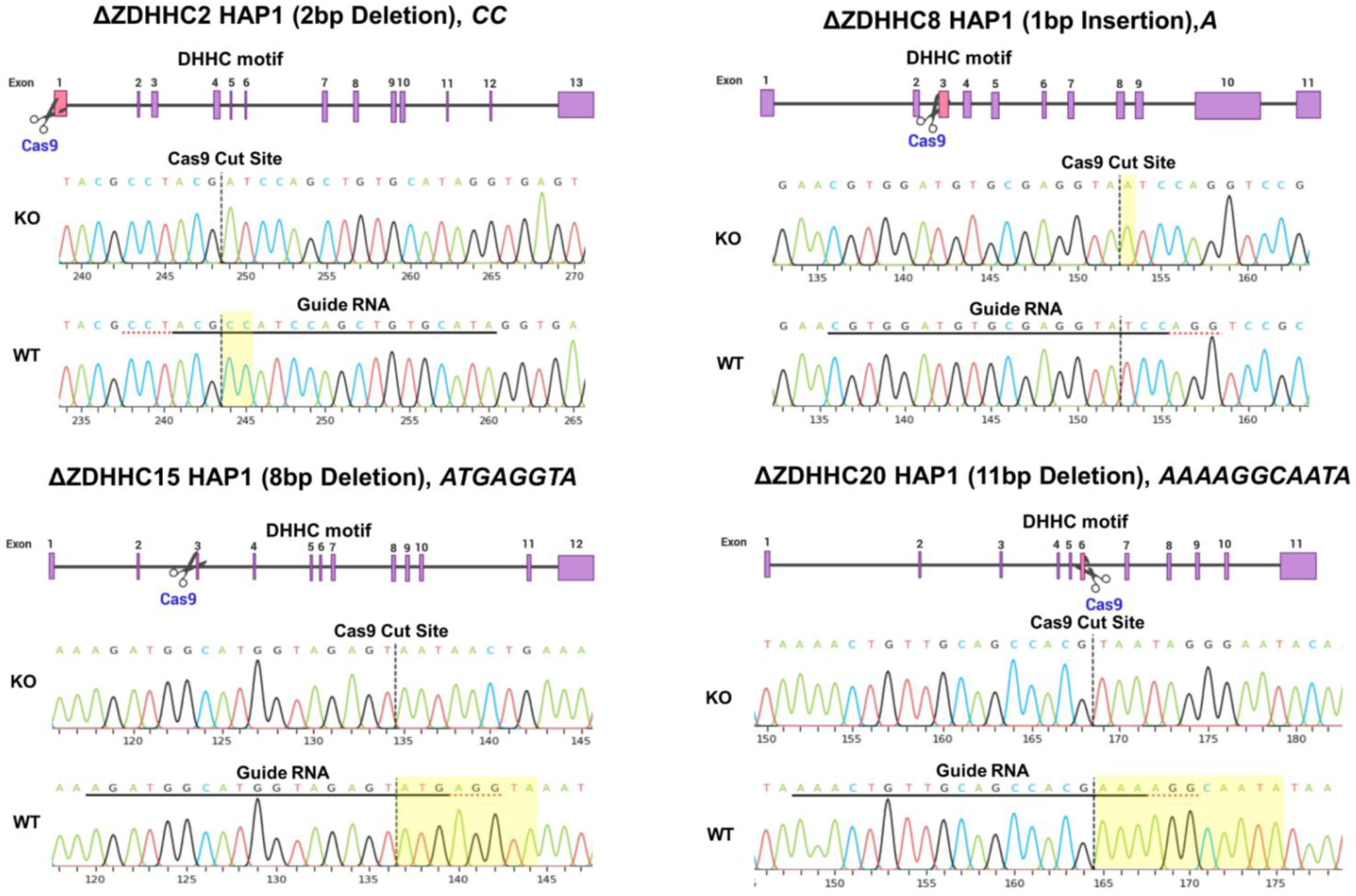
Genotyping of CRISPR/Cas9 induced mutations in ZDHHCs knockout HAP1 cells by PCR and Sanger sequencing. The upper part of each figure shows a scheme of the respective ZDHHC gene. Exons are numbered and marked as purple boxes, the exon which is the target of the guide RNA and hence cleaved by Cas9 is colored pink. The location of the DHHC motif in each ZDHHC gene is also indicated. It is located in exon 6 of ZDHHC2, ZDHHC15 and ZDHHC20 and in exon 4 of ZDHHC8. The lower parts of each figure show two sequencing chromatograms of knockout (KO) and wild type (wt) HAP1 cells. The binding site of the guide RNA is underlined in the sequence. The Cas9 cutting site is indicated by a vertical dotted line. The resulting differences between the WT and KO sequences are highlighted in yellow. The deleted or inserted nucleotides are indicated in the heading of each figure. All induced mutations generate a frameshift in the coding sequence. Parts (330 – 450 bp) of the ZDHHCs gene encompassing the guide RNA binding site were amplified from the genomic DNA by PCR using the primers listed in table S1.

**Figure S5:**
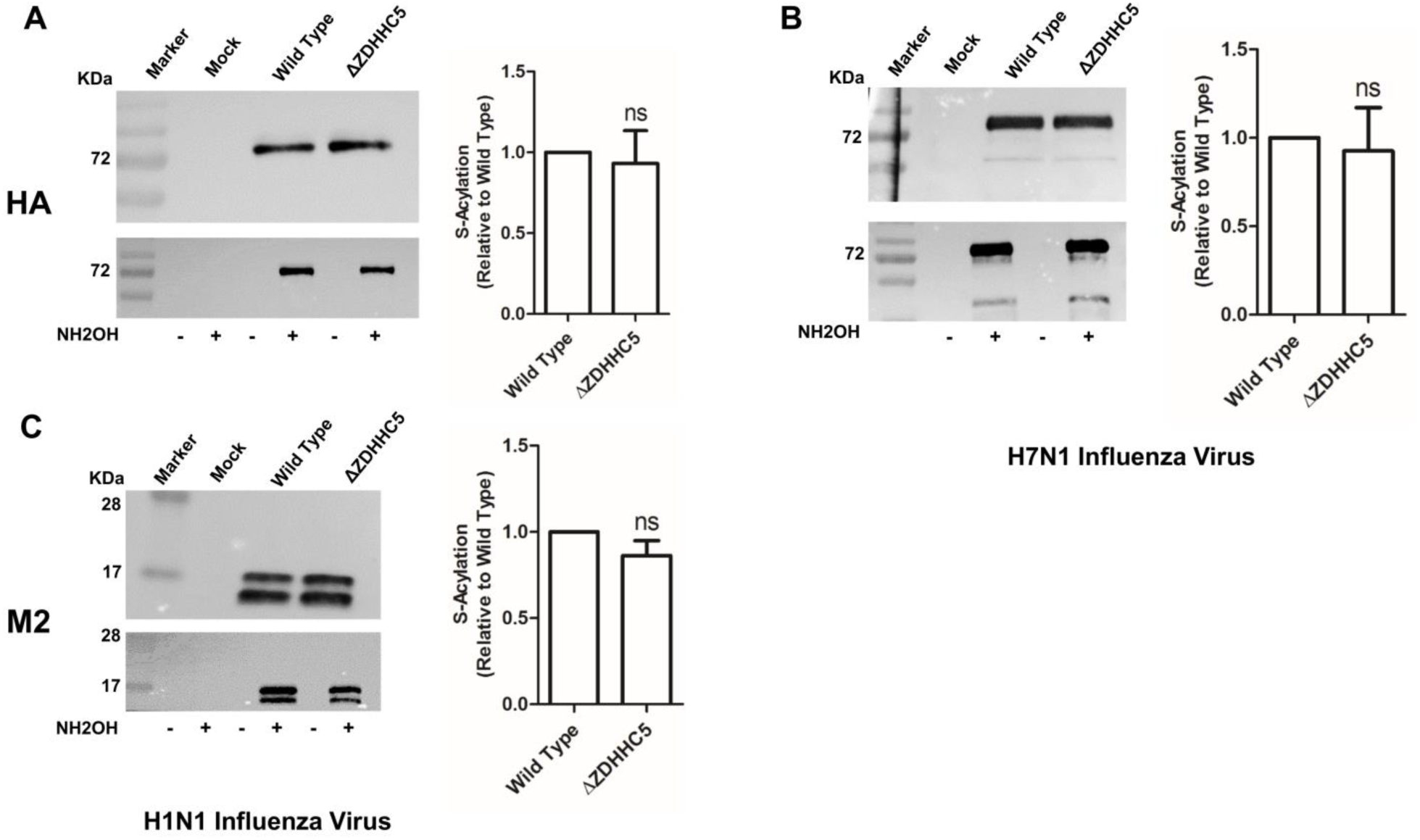
HAP1 cells deficient in ZDHHC5 exhibit no effect on acylation of HA from Flu C and HA from Flu B. **A+C:** ΔZDHHC5 HAP1 cells were infected with the WSN virus (H1N1), which contains a group 1 HA, at an MOI of 1. 24 hours later acylation of HA (A) and M2 (C) proteins were analyzed using Acyl-RAC. B: ΔZDHHC5 HAP1 cells were infected with FPV virus (H7N1), which contains a group 2 HA, at an MOI of 1. 24 hours later acylation of HA was analyzed using Acyl-RAC Quantification of the result from 2 independent experiments is shown. Density of the +NH2OH bands was divided by density of bands from the input and normalized to wild type (=1)). The mean ± standard deviation is shown. One way ANOVA followed by Tukey’s multiple comparison test was applied for statistical analysis. ns: not significant.

**Table S1:**
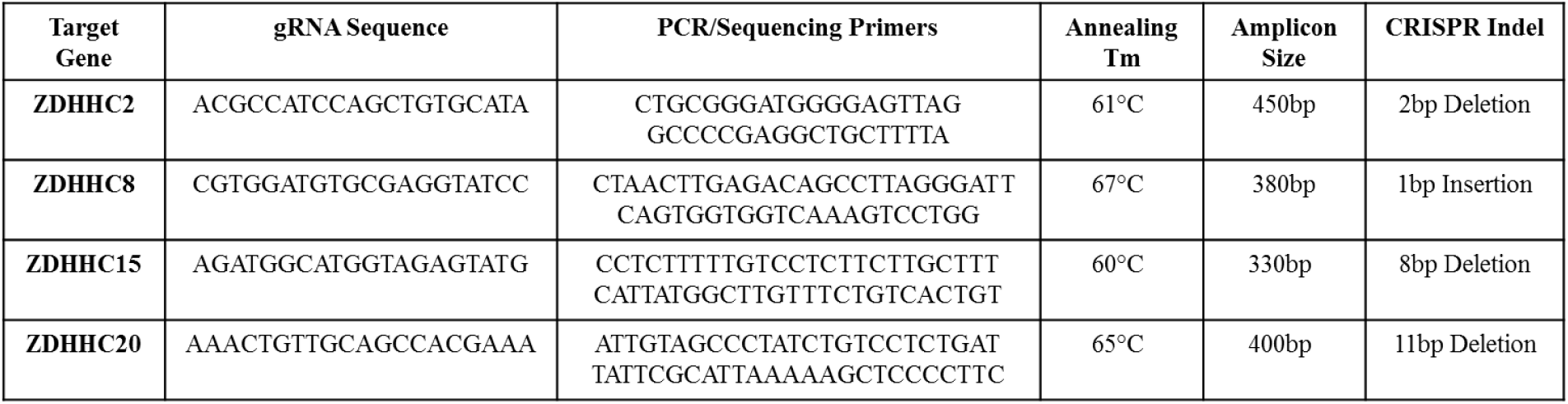
Sequences of guide RNAs and PCR/sequencing primers used for generation and genotyping of CRISPR/Cas9 knockout HAP1 cells. The upper (=forward) of the primers was used for sequencing. The annealing temperature for the PCR reaction, the size in base pairs of the resulting gene fragment and also the number of deleted or inserted nucleotides observed after sequencing of the respective ZDHHC is also listed.

**Table S2:**
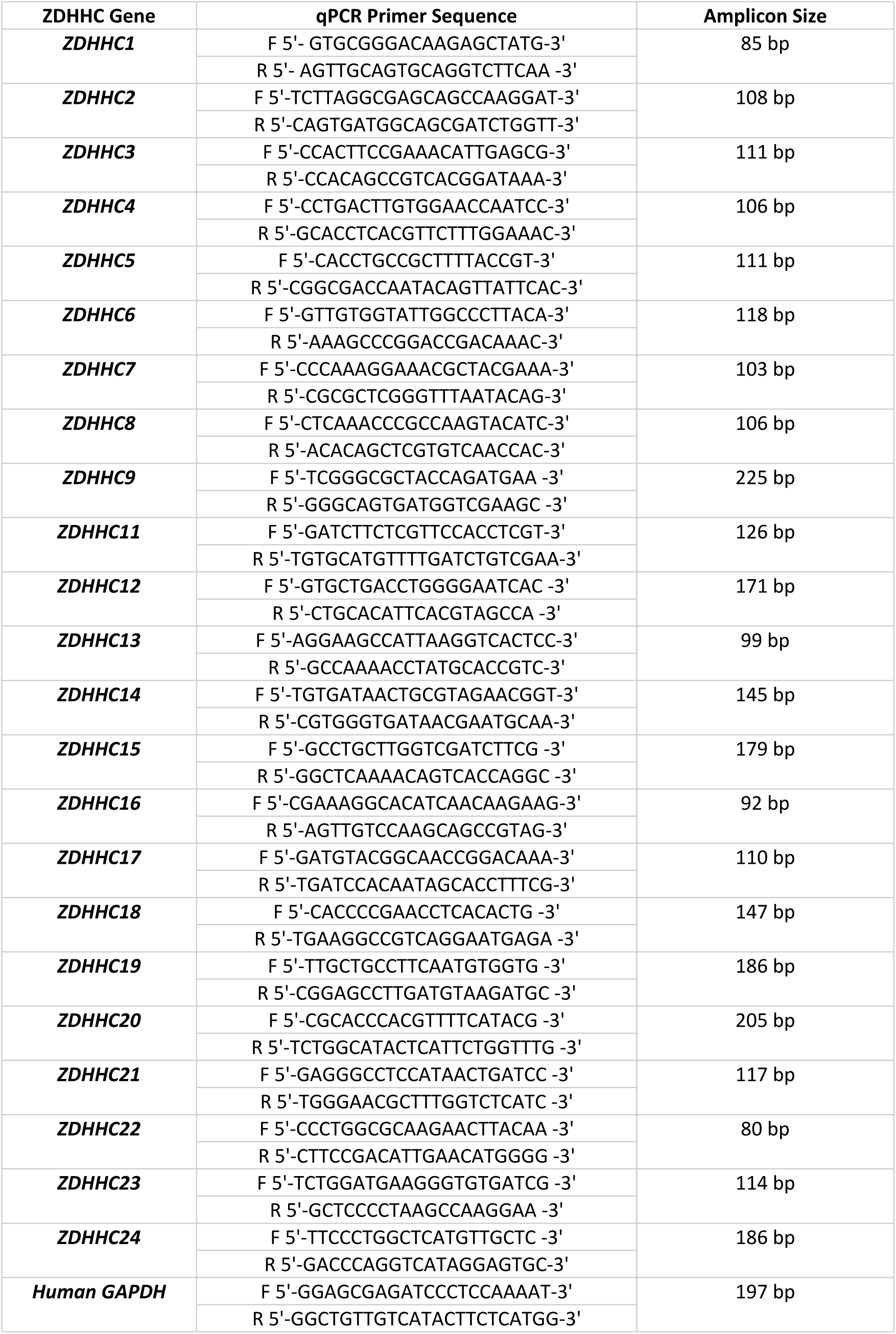
qPCR primers used for analysis of expression of ZDHHCs in human lung cells.

